# MNT loss in MYC-driven B lymphoma cells enhances apoptosis, inhibits proliferation and increases sensitivity to cancer drugs

**DOI:** 10.64898/2026.04.29.721758

**Authors:** Hai Vu Nguyen, Marcel Michla, Feng Yan, Nadia M Davidson, Gemma L Kelly, Andreas Strasser, Suzanne Cory

**Author notes:** Corresponding Author: Suzanne Cory, Walter and Eliza Hall Institute of Medical Research, 1G Royal Parade, Parkville, Victoria 3052, Australia.

## Abstract

Our previous mouse genetic studies showed that loss of the transcription repressor MNT enhanced apoptosis of premalignant lymphoid cells over-expressing MYC and inhibited lymphoma development. Here, we have explored the consequences of inducing *Mnt* deletion in fully malignant lymphoma cells. MNT loss provoked apoptosis of *p53* wt and, albeit more slowly, *p53* mutant *Eμ-Myc* lymphoma cells, preceded by elevated levels of major BH3-only proteins BIM and PUMA. By inhibiting apoptosis, we showed that MNT loss also impaired cell cycling and increased senescence. *Eμ-Myc* lymphoma cells depend on the BCL-2 ortholog MCL-1 for survival and expansion and, importantly, MNT loss enhanced their sensitivity to the MCL-1 inhibitor S63845 and to several chemotherapeutic agents. In BCL-2-overexpressing *Eμ-Myc* lymphoma cells, which model aggressive human ‘Double Hit Lymphomas’, MNT loss enhanced sensitivity to the BCL-2 inhibitor ABT-199, even after BAX loss. Furthermore, *MNT* deletion improved drug responses of two long-established Burkitt Lymphoma cell lines.

**SIGNIFICANCE:** This study establishing the MNT dependency of *Eμ-Myc* lymphoma cells and demonstrating that MNT loss enhances their sensitivity to apoptosis induced by conventional chemotherapeutics and BH3 mimetic drugs provides strong proof-of-concept for developing MNT inhibitors to improve treatment of MYC-driven blood cancers.

## INTRODUCTION

Deregulated overexpression of c-MYC (hereafter MYC) is a major driver of cancer development (1,2). Malignant transformation of MYC-deregulated cells depends on the acquisition of synergistic oncogenic mutations (3,4), but the malignant cells remain MYC-dependent (5). Although much effort has been directed towards developing drugs targeting MYC, a clinically effective drug remains elusive (6,7).

MYC regulates transcription of genes controlling diverse cellular processes, such as cell growth, proliferation, metabolism and the DNA damage response (8). Counter-intuitively, particularly at high expression levels, MYC also induces apoptosis of stressed cells, thereby limiting its oncogenic potential (9–12). Hence, mutations that inhibit apoptosis, such as elevated levels of pro-survival members of the BCL-2 family (13,14), or inactivation of the ARF-MDM2-p53 pathway (15–17), synergize with deregulated MYC in malignant transformation.

MYC is the founding member of a family of related basic Helix-Loop-Helix Leucine Zipper (bHLHLZ) proteins, which includes both gene activators and repressors (for reviews see (8,18,19). MYC heterodimerizes with its obligate partner, MAX, to bind to DNA at canonical and non-canonical CACGTG enhancer box motifs (E-boxes) in target genes. MYC/MAX heterodimers activate transcription of hundreds of genes, although many may be indirect targets (20). Importantly, however, MAX also heterodimerises with MNT and MXD proteins, family members that oppose MYC function by recruiting SIN3 co-repressor complexes to E-boxes to activate histone deacetylases (21).

MNT is widely expressed in mammalian tissues, in both proliferating and quiescent cells, and is vital during development, unlike other MXD family members (8,18,22). As a MYC antagonist, MNT was anticipated to be a tumour suppressor, and this was supported by an early breast cancer study in mice (23). More recently, however, MNT was found to facilitate MYC-driven lymphoma development (24–27), rather than suppressing it. Using *Eμ-Myc* mice (3,4), which model human Burkitt lymphomas driven by chromosome translocations that link *MYC* and *IG* loci (28,29), and two T lymphoma mouse models, we found that *Mnt* deletion greatly reduced lymphoma incidence by enhancing apoptosis of MYC-driven pre-malignant lymphoid cells, at least in part by increasing the level of the BH3-only protein BIM (26,27). BH3-only proteins trigger apoptosis regulated by the BCL-2 protein family. They bind to pro-survival family members such as BCL-2 and MCL-1 and prevent them from inhibiting pro-death relatives BAX and BAK, and certain BH3-only proteins such as BIM and PUMA also directly activate BAX and BAK (30).

Here, we have used *Mnt*-deletable cell lines derived from *Eμ-Myc* lymphomas to further explore early consequences of MNT loss in fully malignant B lymphoid cells. We have also undertaken preclinical studies of *Eμ-Myc* and Burkitt lymphoma cell lines to investigate whether MNT loss enhances sensitivity to conventional cancer drugs and to BH3 mimetic drugs venetoclax and S63845, which activate apoptosis by respectively inhibiting BCL-2 and MCL-1 (31). The results strongly support our proposal (26) that MNT would be a valuable new target for cancer therapy.

## RESULTS

### Inducing *Mnt* Deletion Provokes Apoptosis of *p53* WT and *p53* Mutant *Eμ-Myc* Lymphoma Cells

*Mnt^fl/fl^ Eμ-Myc/Rosa26CreERT2* (hereafter *Mnt^fl/fl^ Eμ-Myc/CreERT2*) mice (26) develop monoclonal pre-B or B lymphomas in which *Mnt* alleles are deletable by exposure to tamoxifen (*in vivo*) or 4-hydroxy tamoxifen (4-OHT) (*in vitro)* (32). We have developed a panel of cell lines (CLs) from *Mnt^fl/fl^* and control *Mnt^+/+^ Eμ-Myc/CreERT2* lymphomas and determined their p53 status by exposure to nutlin3a (33) (Table S1). Here, we have compared the impact of MNT loss in multiple independent *p53* wt and *p53* mutant lymphoma lines using a protocol in which 4-OHT toxicity is minimized by diluting cultures 8-fold at 20 hr with fresh medium lacking 4-OHT (**Fig. 1A**). Under these conditions, control *Mnt*^+/+^ *Eμ*-*Myc/CreERT2* lymphoma CLs (eg #1591; grey curve in **Fig. 1B**) rapidly recovered viability after d2. In contrast, *Mnt^fl/fl^* Eμ-*Myc/CreERT2* lymphoma CLs continued to die, although the *p53* wt (orange) lines died more rapidly than the *p53* mutant lines (red). By d4, *Mnt* deletion was marked for both genotypes (Fig 1C, and not shown). Cell death occurred by the mitochondrial apoptosis pathway, as evidenced by cell surface annexin-V expression and, importantly, BAX/BAK dependency (**Supplementary Figs. S1A-D)**. Of note, *Mnt* deletion did not alter the level of MCL-1, the BCL-2 relative essential for survival of *Eμ-Myc* lymphoma cells (34) (eg **Fig. 1D**).

**Figure 1.**
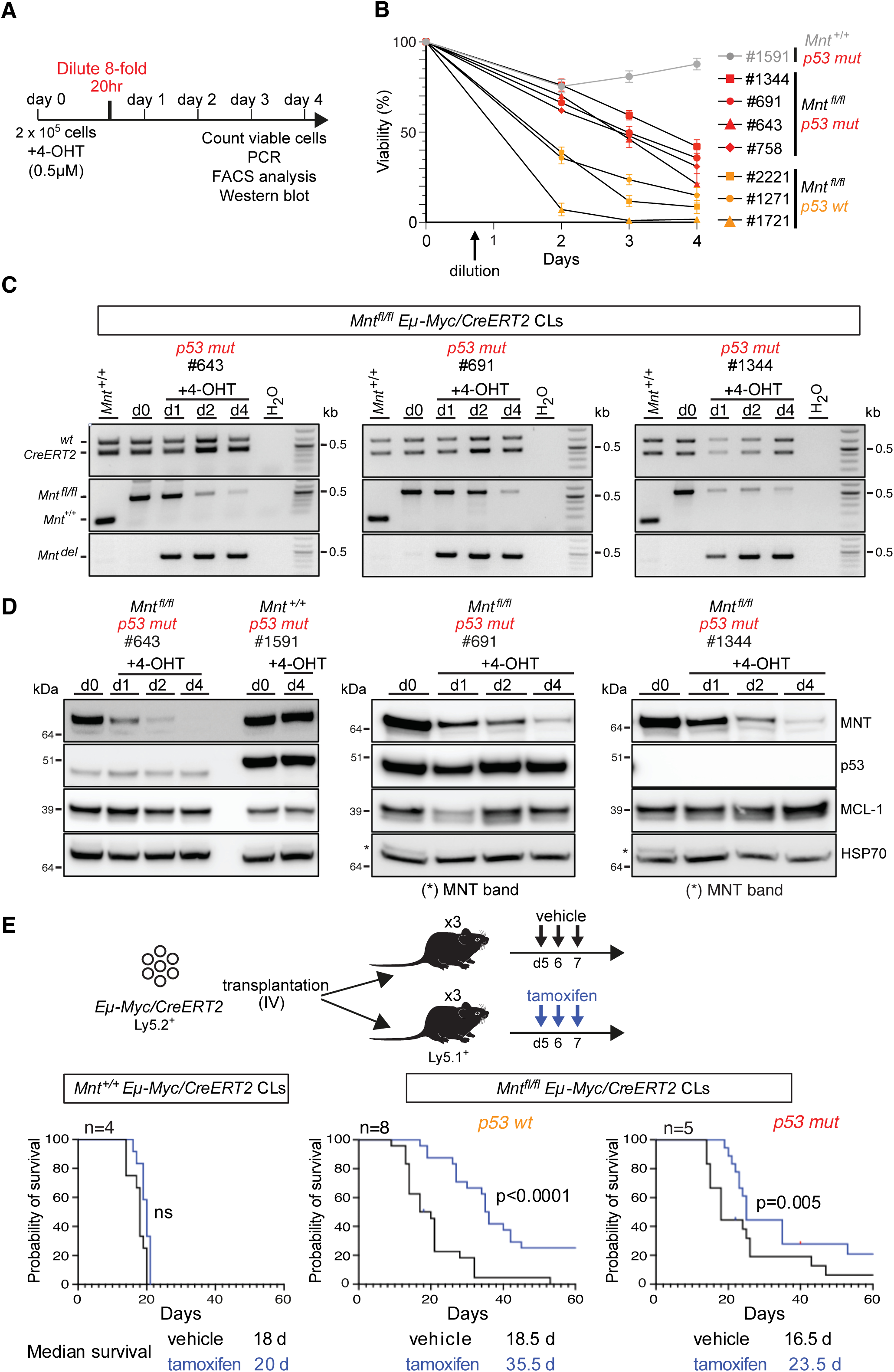
Acute *Mnt* deletion provokes apoptosis of *p53 wt* and *p53 mut Eμ-Myc* lymphoma cell lines (CLs) and extends survival of mice transplanted with these lymphoma cells. **A**, Schematic of protocol used to delete *Mnt* from *Mnt^fl/fl^ Eμ-Myc/CreERT2* lymphoma CLs by brief exposure to 4-OHT (see text). **B**, Viability of 4-OHT-treated *p53 wt* (yellow symbols) and *p53 mut* (red symbols) *Mnt^fl/fl^* Eμ-*Myc/CreERT2* lymphoma CLs compared to that of a control (*p53 mut*) *Mnt^+/+^*Eμ-*Myc/CreERT2* CL (grey symbols), determined by flow cytometry after staining with PI on days 2, 3 and 4. **C**, PCR analysis showing kinetics of deletion of floxed *Mnt* allels in the indicated 4-OHT-treated *p53 mut Mnt^fl/fl^ Eμ-Myc/CreERT2* lymphoma CLs and, as a negative control, *Mnt*^+/+^ *Eμ-Myc/CreERT2* CL #1591. PCR products from *Mnt*^+/+^, *Mnt^fl/fl^* and *Mnt-*deleted (*Mnt^del^*) alleles are indicated, as are products from *Rosa* and *RosaCreERT2* alleles. Size markers are indicated in kb. **D**, Western blot analysis showing kinetics of MNT protein loss after 4-OHT treatment of the indicated *p53 mut Mnt^fl/fl^ Eμ-Myc/CreERT2* lymphoma CLs, ascertained on days 0, 1, 2 and 4. Expression levels of p53 and MCL-1 proteins are also indicated, with HSP70 used as the loading control. Note that the *p53* mutations differ between cell lines: #1591 and #691 express high levels of full length p53 polypeptide, # 643 has lower levels of a smaller polypeptide and #1344, presumably a homozygous deletion, expresses no detectable p53 protein. Asterisk indicates MNT band left from previous immunoblot of this membrane. Protein molecular weight markers are indicated in kDa. **E**, Tamoxifen-induced *Mnt* deletion extends the survival of mice transplanted with *p53 wt* and *p53 mut Mnt^fl/fl^* Eμ-*Myc/CreERT2* lymphoma CLs but not of those transplanted with negative control *Mnt*^+/+^ *Eμ-Myc/CreERT2* lymphoma CLs (see Table S1). Each (Ly5.2^+^) CL was injected into tail veins of 6 non-irradiated (Ly5.1^+^) C57BL/6 recipients (2x10^6^ cells/mouse), 3 of which were treated by oral gavage with tamoxifen and 3 with vehicle (peanut oil containing 10% ethanol) for 3 successive days, starting on day 5; n indicates number of independent lymphoma CLs transplanted. Kaplan-Meier survival curves are depicted, with median survival indicated below; significance was determined by log-rank (Mantel-Cox) test.

To assess the impact of *Mnt* deletion *in vivo*, the (Ly5.2^+^) lymphoma CLs were injected iv into *Mnt^+/+^* unirradiated (Ly5.1^+^) C57BL/6 mice and, after 5 days, the recipient mice were treated daily for 3 days with either tamoxifen or vehicle (3 mice/CL/treatment) (**Fig. 1E**). Mice transplanted with *Mnt^fl/fl^ Eμ-Myc/CreERT2* lymphoma cells survived longer after tamoxifen treatment than those treated with carrier alone, presumably because tamoxifen-induced *Mnt* deletion had provoked substantial apoptosis of the transplanted lymphoma cells. Furthermore, recipient mouse survival significantly improved, irrespective of whether the transplanted CL was *p53* wt or *p53* mutant, although it was greater for the former (median ∼17d versus ∼7d respectively). Tamoxifen-treated control mice transplanted with *Mnt*^+/+^ *Eμ-Myc*/*CreERT2* lymphoma CLs had no survival advantage (left panel).

To assess why transplanted tamoxifen-treated mice eventually relapsed, we analyzed eight tumors that had developed in mice transplanted with three different p53 mutant *Mnt^fl/fl^ Eμ-Myc/CreERT2* lymphomas. Donor-derived lymphoma cells (Ly5.2^+^ CD19^+^) were isolated by FACS from enlarged lymphoid tissues of autopsied mice and analyzed by western blot and PCR (Supplementary Fig. S1E). Some of the tumors (5/8) were positive for MNT protein, indicative of incomplete *Mnt* deletion. In one case (#10337) this was due to deletion of the *CreERT2* transgene, but three other tumors were largely MNT-negative, suggestive of expansion from an apoptosis-resistant sub-population. Consistent with this hypothesis, these tumors (#10301, # 10302, derived from CL #2301, and #10338 derived from CL #2297) had higher levels of pro-survival proteins BCL-2 and/or BCL-X_L_ than those which grew in control mice transplanted with the same parental CLs but treated with vehicle alone.

### BIM and PUMA Levels Rise After *Mnt* Deletion, in Both *p53* WT and *p53* Mutant Eμ- *Myc* Lymphoma Cells

To better understand the upstream signals leading to apoptosis after MNT loss, we overexpressed MCL-1 to prevent the death of *Mnt* ko cells. Multiple independent *Mnt^fl/fl^ Eμ-Myc/CreERT2* lymphoma CLs were infected with *Flag-hMCL-1/GFP* (hereafter *hMCL-1/GFP*) or control *GFP* retroviruses (35) and GFP-positive cells selected by FACS to establish GFP^+^ sub-lines (**Fig. 2A**). As expected, high MCL-1 (MCL-1^hi^) enabled both *p53* wt and *p53* mutant *Eμ-Myc/CreERT2* lymphoma cells to survive in the absence of MNT. Even by d8, their viability was comparable to that of untreated cells (Supplementary Figs. S2A, B).

**Figure 2.**
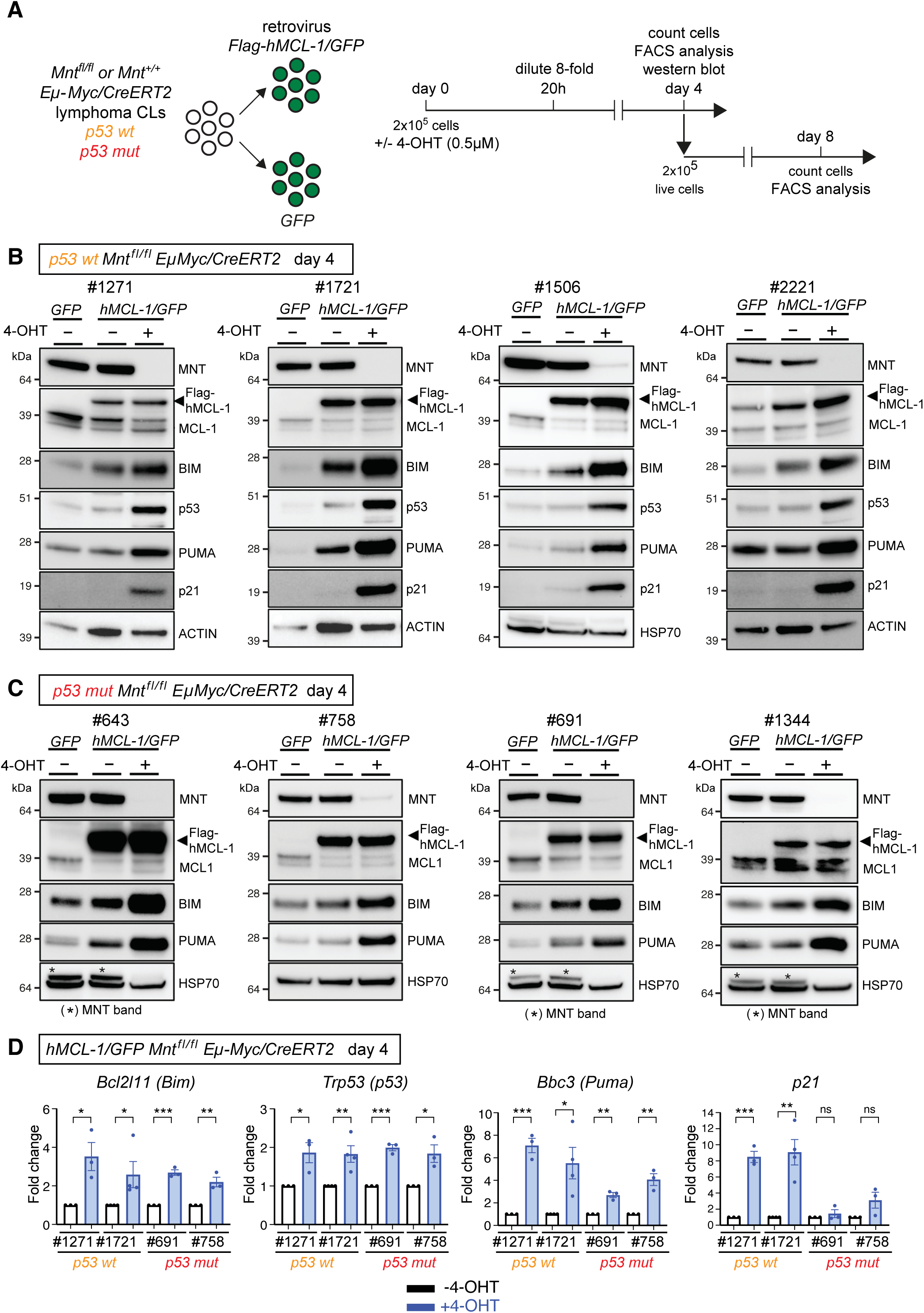
BIM, PUMA and p53 expression is elevated by *Mnt* deletion in *Eμ-Myc* lymphoma cells**. A**, Cartoon summarizing strategy used to identify apoptosis signals upregulated after *Mnt* deletion by deploying MCL-1 over-expressing (*MCL-1^hi^*) *Eμ-Myc* lymphoma cells, which are apoptosis-resistant (see Supplementary Fig. S2A, B). *Mnt^fl/fl^ Eμ-Myc/CreERT2* lymphoma CLs were infected with a *Flag-hMCL-1/GFP* (hereafter *hMCL-1/GFP*) retrovirus or, as a negative control, with a *GFP* retrovirus (see Methods), followed by FACS sorting and culture of GFP-positive cells. The GFP^+^ lymphoma sub-lines were incubated in medium +/- 4-OHT for 20 hr prior to 8-fold dilution with fresh medium and then cultured for subsequent analysis. **B-C,** BIM, PUMA and p53 proteins increase following *Mnt* deletion, in both *p53 wt* (**B**) and *p53 mut* (**C**) *Eμ-Myc* lymphoma cells overexpressing MCL-1, but p21 protein is increased only in *p53 wt* cells. Western blot analysis was performed on day 4 after 20-hr treatment with 4-OHT. Blots are typical of 2 or more independent experiments using different lysates for each lymphoma CL. The viral MCL-1 protein has a higher MW than endogenous MCL-1 due to its N-terminal FLAG tag. The asterisk indicates MNT band left from previous immunoblot performed with the same filter. **D** *Bim*, *Puma* and *p53* mRNAs are elevated after 4-OHT-mediated *Mnt* deletion in independent *p53 wt MCL-1^hi^ Eμ-Myc/CreERT2* lymphoma CLs and *p53 mut MCL-1^hi^ Eμ-Myc* lymphoma CLs. *p21* mRNA is elevated after *Mnt* deletion in *p53 wt* but not *p53 mut* CLs. mRNA levels were determined by Taq-Man PCR analysis on day 4 after the CLs were briefly treated with 4-OHT or medium alone. The bar graphs show fold change of mRNA levels after normalization of 4-OHT treated (blue columns) to medium alone (open columns). Results are shown for at least 3 independent experiments (dots) and expressed as mean ± SEM, normalized to 4-OHT-untreated cells; *P≤ 0.05, **P≤ 0.01, *** P≤ 0.001; ns=not significant.

Western blot analysis (Figs. 2B, C) showed that MNT loss in both the *p53* wt and *p53* mutant CLs was accompanied by increased expression of the BH3-only protein BIM, which is a major trigger of the mitochondrial apoptosis pathway regulated by the BCL-2 family (30), *Bim* transcription was also significantly elevated (panel 1, Fig. 2D).

In the *p53* wt MNT-deficient lymphoma cells, p53 protein and mRNA also increased (Figs. 2B, D), accompanied by increased expression of p53 target genes: *Bbc*, which encodes PUMA, another pro-apoptotic BH3-only protein, and *Cdkn1A*, which encodes the cell cycle inhibitor p21 (36). It might be argued that CRE-mediated DNA cleavage could be responsible for the increased p53. To directly address this possibility, we investigated the kinetics of *Mnt* deletion and p53 expression (Supplementary Fig. S2D). Treatment with 4-OHT was terminated at 20 hr and Cre-mediated *Mnt* deletion was well-advanced when analyzed at 24 hr. Increased expression of p53, PUMA and p21 protein was not apparent until 48 hr and remained high at 96 hr, when CRE activity would be expected to be negligible. Furthermore, elevated expression of p53, PUMA and p21 was still apparent in all *p53* wt *Mnt-ko* subclones at 4 weeks, long after any CRE activity could remain (Supplementary Fig. S2E). These results argue against CRE-mediated DNA breaks as the cause of the p53 increase and instead suggest that *Mnt* deletion has activated *p53* expression.

In the *p53* mutant *MCL-1^hi^* CLs, PUMA protein and *Bbc* (*Puma*) RNA also increased after *Mnt* deletion (Figs. 2C, D) but p21 was not detectable (Supplementary S2F). Thus, p53-independent mechanisms also increase PUMA levels after *Mnt* deletion. As expected, BIM and PUMA levels remained unaffected in control *Mnt*^+/+^ *p53* mutant cells treated with 4-OHT (Supplementary Fig. S2C).

Pertinently, analysis of published CUT&RUN data from wt and *Max^-/-^*B220^+^ splenocytes (37) suggest that *Bim* (*Bcl211*), *Puma (Bbc3)* and *p53 (Trp53)* genes are all direct targets of MNT/MAX heterodimers, as is *Mnt* itself (Supplementary Fig. S2G).

Taken together, these data imply that MNT-mediated suppression of pro-apoptotic genes *Bim*, *Puma* and *p53* protects MYC-driven lymphoma cells from excessive apoptosis.

### MNT Loss in Apoptosis-Resistant *Eμ-Myc* Lymphoma Cells Results in Reduced Cell Proliferation and Increased Senescence

The apoptosis-resistant *MCL-1^hi^ Mnt*-deletable *Eμ-Myc* lymphoma CLs enabled us to also assess the impact of MNT loss on cell proliferation. Despite retaining high viability after 4-OHT treatment (Supplementary Fig. S2A, B), 4-OHT treated CLs grew very slowly compared to untreated controls, irrespective of their *p53* status (compare blue with open columns in Fig. 3A).

**Figure 3.**
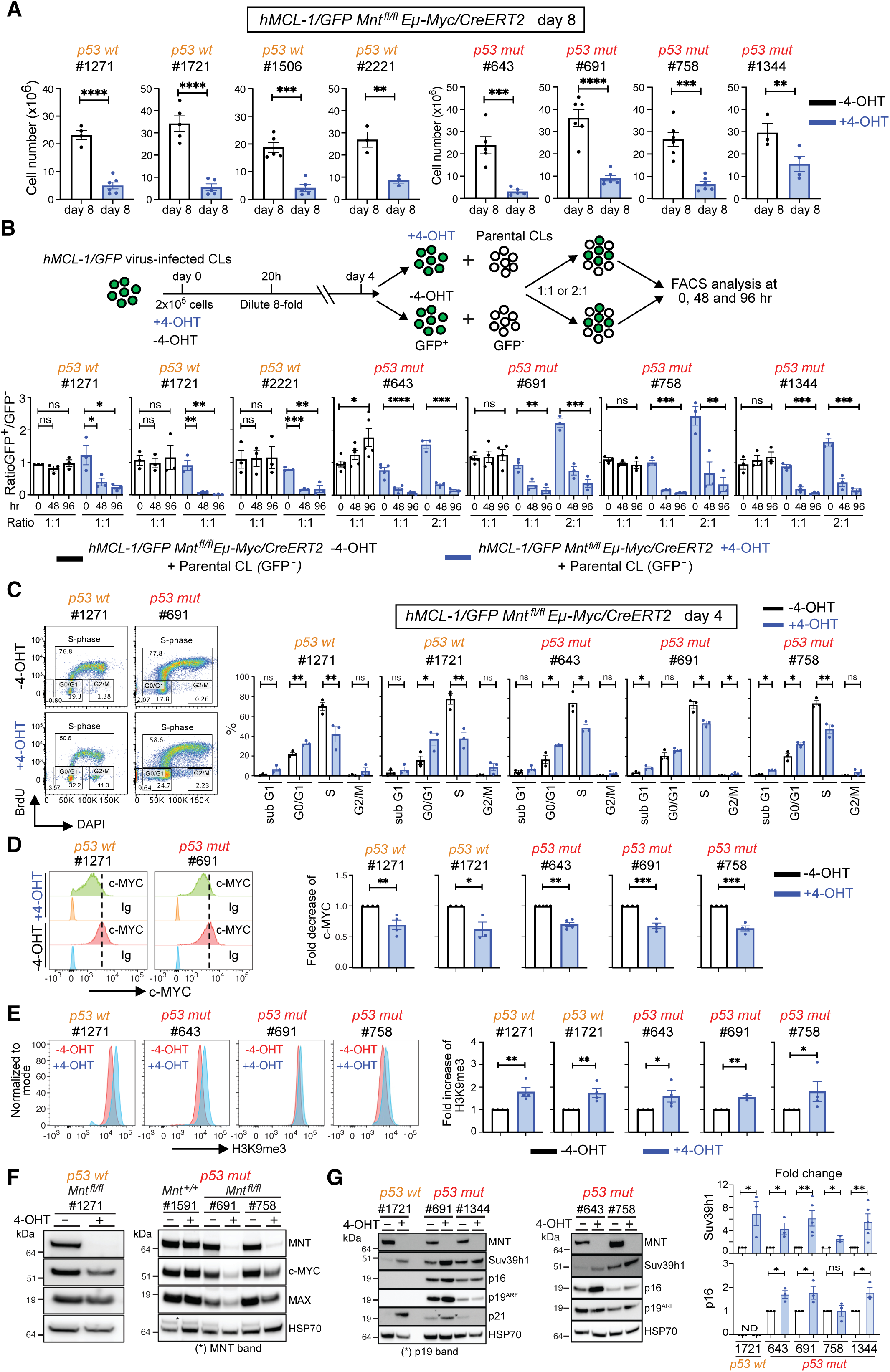
MNT loss in apoptosis-resistant (*MCL-1^hi^*) *Eμ-Myc* lymphoma cells reduces cell proliferation and increases cell senescence. **A**, Reduced proliferation of *MCL-1^hi^ p53* wt and *p5*3 *mut Mnt^fl/fl^ Eμ-Myc/CreERT2* lymphoma CLs following 4-OHT-induced *Mnt* deletion. Cells were treated for 20 hr +/- 4-OHT then diluted and cultured as shown in Fig 2A. Cell number on day 8 is compared for cells treated with 4-OHT (blue columns) or medium alone (open columns). **B**, Competition assay showing reduced proliferation of *Mnt*-deleted *MCL-1^hi^ p53 wt* and *p53 mut Eμ-Myc* lymphoma cells. Indicated *MCL-1^hi^ Mnt^fl/fl^ Eμ-Myc* lymphoma CLs (GFP^+^) were treated briefly +/- 4-OHT, harvested on day 4 and then competed 1:1 or 2:1 against uninfected parental CLs (GFP^-^). The relative proportion of GFP^+^/GFP^-^ cells was determined by FACS at 0, 48 and 96 hr, as exemplified in Fig S3A. The protocol is outlined in the upper panel and bar graphs below show diminishing GFP^+^/GFP^-^ ratios over time. Three experiments were performed for each indicated independent *p53* wt and *p53* mut *Mnt^fl/fl^ Eμ-Myc/CreERT2* lymphoma CL, and for a control *Mnt^+/+^* (#1591) cell line (see Fig S3B). **C**, Cell cycling is reduced after *Mnt* deletion. After brief treatment +/- 4-OHT, *MCL-1^hi^ Mnt^fl/fl^ Eμ-Myc/CreERT2* CLs were labelled with BrdU for 3hr on day 4, then stained with BrdU antibody and DAPI for cell cycle analysis by flow cytometry (see Methods). A typical analysis is shown on the LHS and bar graphs summarise % of cells in different stages of the cycle for cells pre-treated with 4-OHT (blue columns) or medium alone (open columns). **D**, MYC protein levels are reduced after *Mnt* deletion. After brief treatment +/- 4-OHT, permeabilized cells were stained on day 4 with antibodies against MYC or, as a negative control, Ig. MYC level, was determined by flow cytometry, as exemplified in left panel. Bar graphs summarize the fold decrease in median fluorescence of MYC staining in cells treated with 4-OHT (blue columns) versus medium alone (open columns). **E**, Heterochromatin H3K9 trimethylation is increased after *Mnt* deletion. H3K9me3 staining of permeabilized cells of the indicated genotypes on day 4, analyzed by FACS. A typical analysis is shown on the left and bar graphs on the right show fold increase in median fluorescence of H3K9me3 in cells treated with 4-OHT (blue columns) versus medium alone (open columns). **F**, Expression of MNT, MYC and MAX proteins in indicated *Mnt^fl/fl^* and control *Mnt^+/+^Eμ-Myc/CreERT2* lymphoma CLs on day 4 following brief treatment +/- 4-OHT. **G,** Expression of cell cycle inhibitors p16, p19Arf, p21, and histone methyltransferase Suv39h1, which catalyzes trimethylation of H3K9, following *Mnt* deletion in indicated *MCL-1^hi^ Mnt^fl/fl^ Eμ-Myc/CreERT2* lymphoma CLs on day 4 following brief treatment +/- 4-OHT. Expression was determined by western blot and examples shown are typical of at least 3 independent experiments, with HSP70 as a loading control. Asterisks indicate band from previous blot performed with the indicated antibody. The bar graphs show the fold change of Suv39h1 and p16 protein in the indicated *MCL^hi^ Eμ-Myc* lymphoma CLs in cells treated with 4-OHT (blue columns) versus medium alone (open columns), determined by quantification of the specific signal relative to the HSP70 loading control. ND = undetermined.

Cell competition assays provided further evidence for the cell proliferation defect (Fig. 3B, Supplementary Fig. S3A). Following brief 4-OHT-treatment, *hMCL-1/GFP* retrovirus-infected *Mnt^fl/fl^ Eμ-Myc/CreERT2* lymphoma cells harvested on d4 were mixed 1:1 or 2:1 with untreated, uninfected (GFP^-^) parental cells and their relative proportions assessed by flow cytometry after 0, 48 and 96 hr in culture. In the cultures containing 4-OHT-treated *Mnt^fl/fl^* lymphoma CLs (blue columns), the proportion of GFP^+^ cells diminished over time, for both *p53* wt or *p53* mutant CLs, whereas the ratio remained constant for controls treated with medium alone (open columns). Note that treatment with 4-OHT did not affect the proliferation of control *MCL-1^hi^ Mnt^+/+^ Eμ-Myc/CreERT2* lymphoma cells (Supplementary Fig. S3B).

Cell competition assays were also performed with multiple *Mnt ko* clones isolated by single cell cloning from a *p53* wt and a *p53* mutant *hMCL-1/GFP* virus-infected *Mnt^fl/fl^ Eμ-Myc/CreERT2* CLs 8 days after 4-OHT treatment (Supplementary Fig. S3C). In each case, the (GFP^+^) *Mnt*^-/-^ clones grew more slowly than the (GFP^-^) parental CL.

Finally, cell competition assays were performed with (BFP^+^) *BaxBak dko Mnt^fl/fl^ Eμ-Myc/CreERT2* lymphoma CLs isolated by CRISPR/Cas9 (Supplementary Fig. S1D). Two independent clones of one *p53* wt and three *p53* mutant CLs were pre-treated with or without 4-OHT and then cultured 1:1 against the parental CL (BFP^-^) for 4 days (Supplementary Fig. S3D). In every case, irrespective of p53 status, 4-OHT-treated (+) (ie *Mnt*-deleted) cells competed poorly compared to untreated cells (-).

Thus, when the mitochondrial apoptosis pathway was blocked, either by elevated MCL-1 or loss of both BAX and BAK, MNT-deprived lymphoma cells survived well but proliferated poorly.

Cell cycle analysis of BrdU-labelled cells confirmed the reduced proliferation (Fig. 3C). Whether *p53* wt or *p53* mutant, MNT-depleted (4-OHT-treated) *MCL-1^hi^ Eμ-Myc* lymphoma cell populations (blue columns) showed a significant reduction in S-phase and an increase in G0/G1 phase cells compared to control untreated cells (open columns).

Reduced MYC protein levels within the 4-OHT-treated *MCL-1^hi^ Mnt^fl/fl^ Eμ-Myc/CreERT2* CLs (Figs. 3D, F) and derivative *Mnt ko* clones (Supplementary Fig. S3E), together with elevated H3K9me3 (Fig. 3E) and histone methyl transferase Suv39h1 (Fig. 3G), plus expression of cell cycle inhibitors p21 (in *p53* wt CLs; Fig. 2B) and p16 (in *p53* mutant CLs; Fig. 3G), all suggested that cells were accumulating in a senescent state (38). The GFP expressed by *MCL-1^hi^ Eμ-Myc* lymphoma cell populations precluded assessment of SA- β -Gal activity as a marker of senescence. However, using the *BaxBak dko* lymphoma clones, which are BFP^+^, we were able to detect an increase in SA- β -Gal-positive cells (Supplementary Fig. S3F).

In summary, MNT-deficient *Eμ-Myc* lymphoma cells that have acquired apoptosis resistance have poor proliferative capacity. These results have important therapeutic implications (see Discussion).

### *Mnt* Deletion Increases Sensitivity to Conventional Cancer Therapeutics

To assess whether MNT loss increased sensitivity to conventional anti-cancer drugs, we tested three agents that induce apoptosis and are commonly used in the clinic to treat lymphoma patients: Cisplatin, which cross links DNA, inhibiting DNA replication and transcription; Doxorubicin, which intercalates into DNA and disrupts topoisomerase-mediated DNA repair; and Vincristine, which inhibits microtubule aggregation, thereby blocking cell division (39). Independent *p53* wt and *p53* mutant *Mnt^fl/fl^ Eμ-Myc/CreERT2* lymphoma CLs infected with the *hMCL-1/GFP* or control *GFP* retrovirus were exposed briefly (or not) to 4-OHT on d1 to delete *Mnt*, then treated on d4 for 24 hr with a range of concentrations of the chemotherapeutic drug (Supplementary Fig. S4A).

Sensitivity to Cisplatin (Fig. 4A) was high for MNT-positive (4-OHT untreated) GFP-virus-infected *p53* wt CLs and, as expected, resistance was increased by *p53* mutations (compare black curves in top and bottom panels). High levels of MCL-1 increased the resistance of both *p53* wt and *p53* mutant CLs (compare black and red curves in top and bottom panels). However, *Mnt* deletion (4-OHT treatment), which increased BIM and PUMA levels (Fig 2B,C and Supplementary Fig S4B), significantly increased the Cisplatin sensitivity of all three testable *GFP*-virus-infected *p53* mutant CLs (#691, #758, #1344) (compare blue to black lines), all h*MCL-1* virus-infected *p53* wt lines (compare green to red curves in top panels) and 2/4 MCL^hi^ *p53* mutant lines (# 691, #1344) (compare green to red curves in bottom panels).

**Figure 4.**
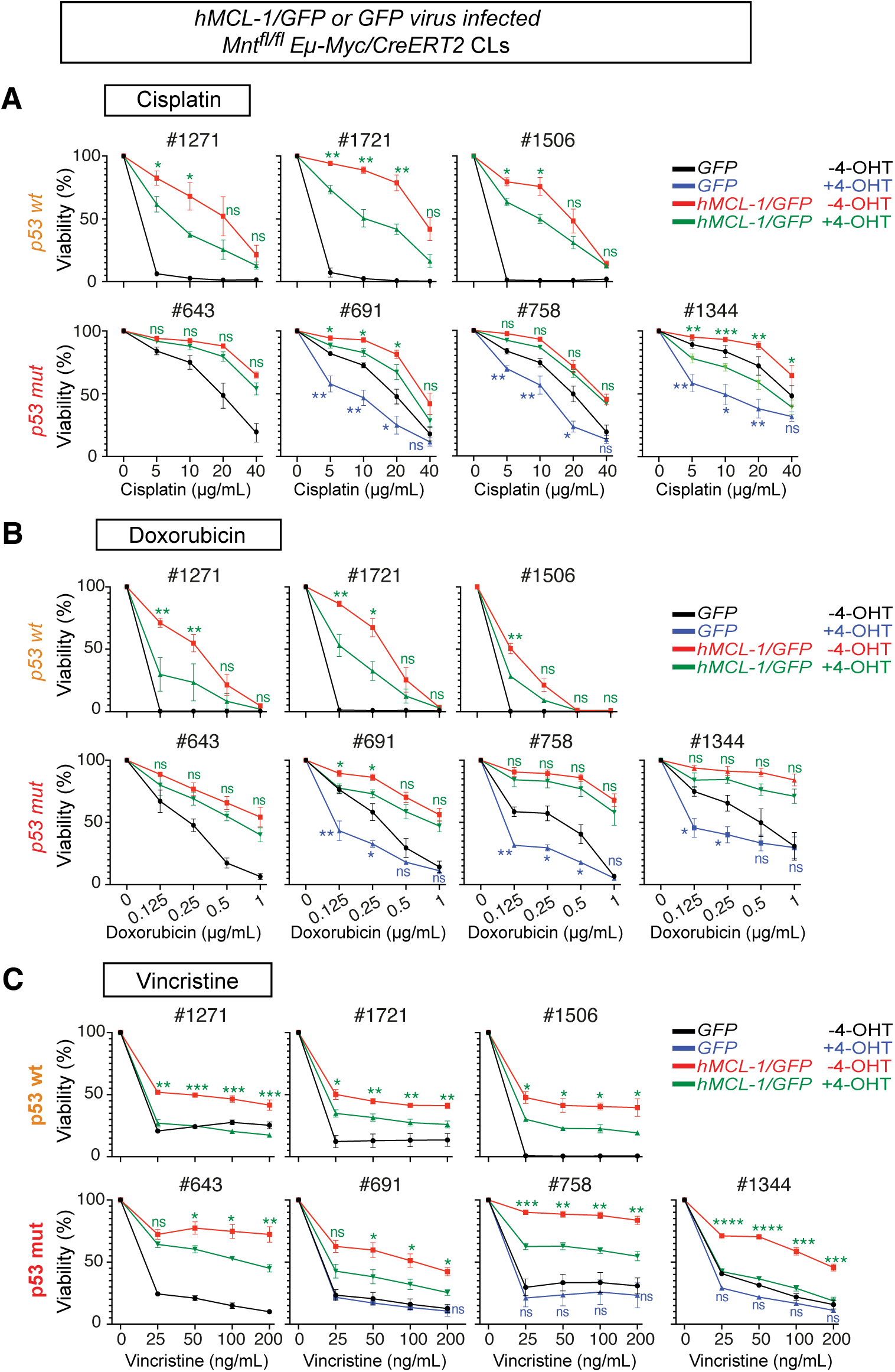
MNT loss increases the sensitivity of *Eμ-Myc* lymphoma cells to cancer chemotherapeutic drugs. **A-C**, Comparison of the sensitivity of *p53 wt* and *p53 mut Mnt ^fl/fl^ Eμ-Myc*/*CreERT2* lymphoma CLs infected with the *hMCL-1/GFP* or control *GFP* retrovirus to Cisplatin, Doxorubicin and Vincristine, with or without prior MNT loss. The indicated CLs were briefly treated (or not) with 4-OHT (0.5μM) followed by dilution as described in **Fig S4A,** then treated on day 4 with a range of concentrations of the indicated drug for 24 hr before determining viability by flow cytometry. The drug sensitivity of *GFP* virus-infected cells pre-treated with 4-OHT or medium alone is indicated as blue and black lines respectively and the drug sensitivity of *hMCL-1/GFP* virus-infected cells pre-treated with 4-OHT or medium alone is indicated in green and red respectively. Due to their very high sensitivity to MNT loss alone, combination therapy (blue lines) could not be examined for the p53 wt CLs infected with GFP virus or *p53* mut line #643 infected with *GFP* virus, but was readily assessed for all *MCL-1/GFP* virus-infected CLs. Green asterisks indicate significance of differences in sensitivity between green (+ 4-OHT) and red (- 4-OHT) curves for *hMCL-1/GFP* retrovirus-infected CLs. Blue asterisks indicate significance of differences in sensitivity between blue (+ 4-OHT) and black (-4-OHT) curves for *GFP* virus-infected CLs. Results are shown for at least 3 independent experiments and presented as mean ± SEM; *P≤ 0.05, **P≤ 0.01, *** P≤ 0.001, ns= not significant.

Likewise, with Doxorubicin (Fig. 4B), sensitivity was high for MNT-positive p53 wt CLs and substantially reduced by p53 mutation (compare black lines between top and bottom panels), and prior *Mnt* deletion in the 3 testable *p53* mutant CLs significantly increased sensitivity (compare blue to black lines in bottom panel). High MCL-1 levels also increased resistance to Doxorubicin and alleviation by *Mnt* deletion was variable between the CLs (compare green to red curves).

With Vincristine (Fig. 4C), p53 mutation had less impact on the sensitivity of the parental lines (compare black curves in top and bottom panels). However, while elevated MCL-1 levels increased the resistance of both *p53* wt and *p53* mutant CLs, prior MNT loss significantly restored sensitivity (compare green to red curves).

### *Mnt* Deletion Increases Sensitivity to BH3 Mimetic Drug S63845

In view of the known MCL-1 dependence of *Eμ-Myc* lymphomas (34), we next tested the BH3 mimetic drug S63845 (40), which binds to MCL-1 and prevents it from inhibiting apoptosis-inducing BAX and BAK (31,41,42). All control (GFP virus-infected) *Eμ-Myc* CLs (black curves in Fig. 5A) were sensitive to the MCL-1 inhibitor, irrespective of *p53* status and overexpression of hMCL-1 (which has a 7-fold higher affinity for S63845 than mouse MCL-1) (40) (red curves) significantly increased resistance. Treatment with 4-OHT to induce *Mnt* deletion improved the S63845-sensitivity of the more resistant *p53* mutant lines #691 and #758 (compare blue to black lines) and significantly enhanced the responsiveness of all *MCL-1^hi^* CLs (compare green to red curves) but had no impact in negative control *Mnt*^+/+^ *Eμ-Myc/CreERT2* CLs (Supplementary Fig. S5A).

**Figure 5.**
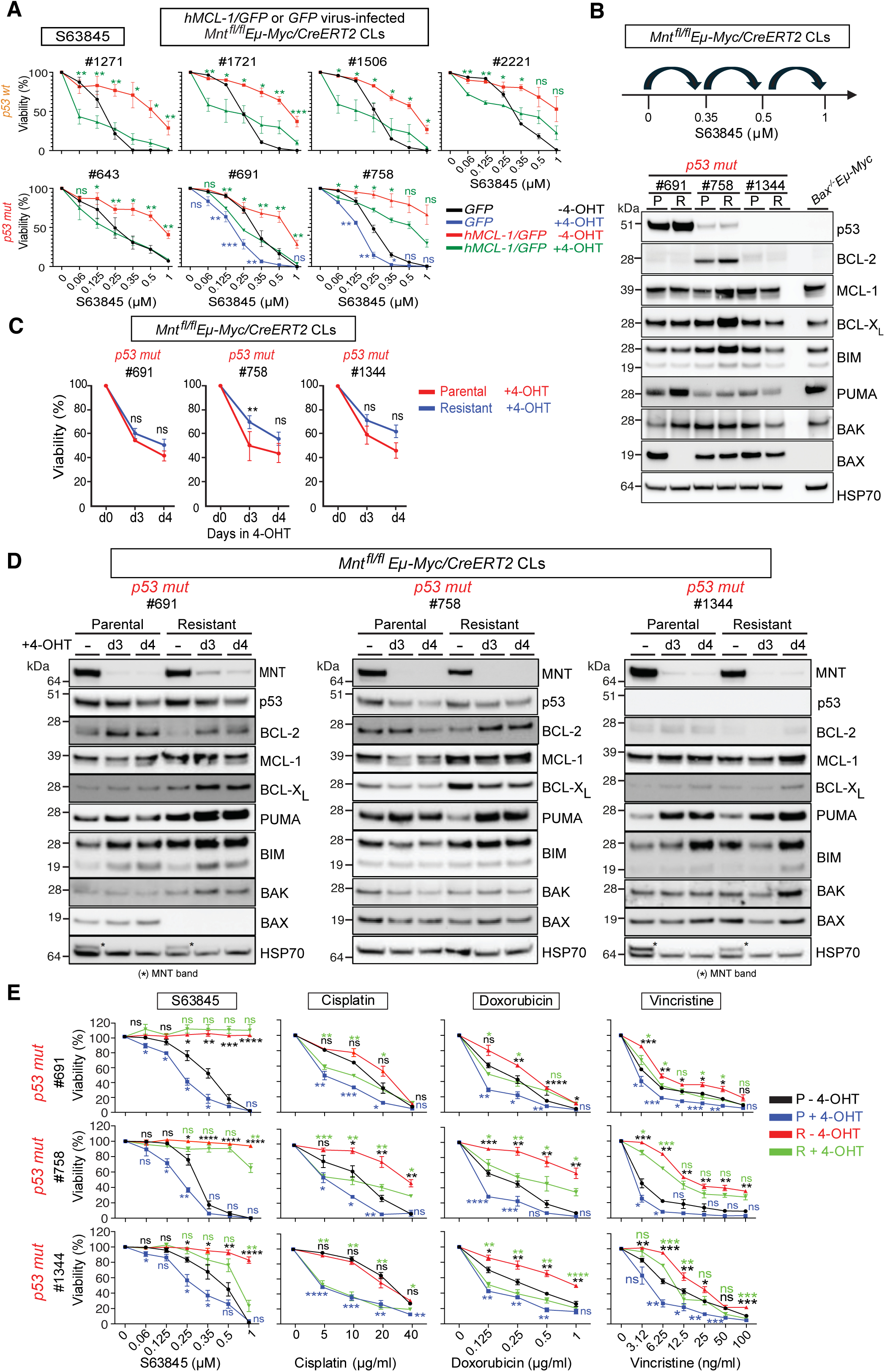
*Mnt* deletion increases the sensitivity of *Eμ-Myc* lymphoma cells to the MCL-1-specific BH3 mimetic S63845 and enhances the response of derivative S63845-resistant cells to Cisplatin, Doxorubicin and Vincristine**. A**, Impact of MNT loss on sensitivity of h*MCL-1*/*GFP* and *GFP* retrovirus-infected *p53 wt* and *p53 mut Mnt^fl/fl^ Eμ-Myc/CreERT2* lymphoma CLs to S63845. Lymphoma cells were treated +/- 4-OHT (0.5μM) followed by dilution as described in Supplementary Fig. S4A, then treated on day 4 with the indicated concentrations of S63845 for 24 hr before determining viability by flow cytometry. Sensitivity of *GFP* virus-infected lymphoma cells that had been exposed to 4-OHT or medium alone is indicated in blue and black respectively. Sensitivity of *hMCL-1/GFP* virus-infected lymphoma cells that had been exposed to 4-OHT or medium alone is indicated in green and red respectively. Green asterisks indicate significance of differences in sensitivity between green (+ 4-OHT) and red (-4-OHT) curves for *hMCL-1/GFP* retrovirus-infected lymphoma cells, and blue asterisks indicate significance of differences in sensitivity between blue (+ 4-OHT) and black (- 4-OHT) curves for *GFP* retrovirus-infected lymphoma cells. Results are shown for at least 3 independent experiments for each indicated CL and plotted as mean ± SEM; *P≤ 0.05, **P≤ 0.01, *** P≤ 0.001, ns= not significant. **B**, Generation and characterization of S63845-resistant *Mnt^fl/fl^ Eμ-Myc/CreERT2* lymphoma CLs. Upper diagram depicts isolation of S63845-resistant lymphoma cells by exposure to increasing concentrations of drug (see Methods). Western blot below shows expression of p53 and indicated BCL-2 family proteins in parental (P) and resistant (R) derivative sub-lines with HSP70 as a loading control. **C,** S63845-resistant *p53 mut Mnt^fl/fl^ Eμ-My/CreERT2* lymphoma cells remain sensitive to MNT loss. Viability of parental (P, red) and S63845-resistant (R, black) lymphoma cells, determined by flow cytometry 0, 3 and 4 days after 4-OHT treatment as described in Fig 1A. **D**, Western blots comparing the level of MNT, p53 and BCL-2 family proteins in the indicated *p53 mut* parental (P) and derivative S63845-resistant (R) lymphoma CLs prior to (-) and on day 3 and day 4 after 4-OHT treatment to delete *Mnt*. **E**, MNT loss enhances sensitivity of *p53 mut* S63845-resistant *Eμ-Myc* lymphoma CLs to Cisplatin, Doxorubicin and Vincristine. Indicated parental and S63845-resistant *Mnt^fl/fl^ Eμ-Myc/CreERT2* lymphoma CLs were treated briefly +/- 4-OHT (0.5 μM) followed by dilution, then treated on day 4 with increasing concentrations of the indicated chemotherapeutic drugs for 24 hr before determining viability by flow cytometry. Curves compare the drug sensitivity of *Mnt* wt (-4-OHT) and *Mnt*-deleted (+4-OHT) parental (P) and S63845-resistant (R) lymphoma cells to S63845, Cisplatin, Doxorubicin and Vincristine. Green asterisks indicate significance of differences in sensitivity between green (+4-OHT) and red (- 4-OHT) curves for S63485-resistant (R) lymphoma CLs treated with the indicated drug. Blue asterisks indicate significance of differences in sensitivity between blue (+4-OHT) and black (-4-OHT) curves for parental (P) lymphoma CLs. Black asterisks indicate significance of differences in sensitivity between 4-OHT-untreated parental (black) and S63845-resistant (red) curves. Results are shown for at least 3 independent experiments for each lymphoma CL, plotted as mean ± SEM; *P≤ 0.05, **P≤ 0.01, *** P≤ 0.001, ns= not significant.

Despite initial sensitivity, blood cell cancers treated with BH3 mimetic drugs can eventually evolve resistance eg (43–46). To test whether MNT loss could still increase the sensitivity of S63845-resistant lymphoma cells to conventional drugs, we derived sublines resistant to >1μM of S63845 from three *p53* mutant and two *p53* wt *Mnt^fl/fl^ Eμ-Myc/CreERT2* CLs (Fig. 5B and Supplementary Fig. S5B, C). Western blot analysis of the *p53* mutant CLs (#691, #758, #1344) suggested that loss of BAX expression is responsible for the acquired resistance of sub-line #691^R^ (compare first 2 tracks of Fig 5B). For CL #758, the resistance of sub-line #758^R^ is probably due to the increased level of MCL-1, BCL-2 and BCL-X_L_, but the basis for the resistance developed by sub-line #1344^R^ remains unclear. For the S63845-resistant *p53* wt CLs (Supplementary Fig. S5C), an increased BCL-X_L_ level could account for the resistance of the #1271^R^ subline, but it is unclear why #1721^R^ is more resistant. These results confirm and extend a previous study of *p53* wt *Eμ-Myc* CLs selected for S63845-resistance (47).

Importantly, the S63845-resistant CLs still died in response to loss of MNT (Fig. 5C and Supplementary Fig. S5D) which, as in the S63845-sensitive parental cells, was accompanied by increased levels of BIM and PUMA (Fig. 5D).

Therefore, we tested whether MNT loss could enhance the sensitivity of the S63845-resistant *p53* mutant CLs to traditional chemotherapeutics (Fig. 5E). Each of the S63845-resistant CLs was more resistant to doxorubicin and vincristine than the parental cells but only CL #758 was also more resistant to Cisplatin (compare red and black curves). Importantly, 4-OHT-induced MNT loss significantly increased the sensitivity of the S63845-resistant CLs to cisplatin and doxorubicin, and, to a lesser extent, to vincristine (compare green to red curves).

Taken together, these results suggest that targeting MNT would be an effective strategy for enhancing the sensitivity of many *p53* wt and *p53* mutant MYC-driven lymphomas to diverse anti-cancer drugs, including the BH3 mimetic S63845.

### *Mnt* Deletion Enhances Venetoclax Sensitivity of MYC-driven Lymphoma Cells Overexpressing BCL-2

In contrast to their sensitivity to S63845, *Eμ-Myc* lymphoma cells are very resistant to BH3 mimetics ABT-199 (venetoclax), a specific BCL-2 inhibitor (48,49), and ABT-737 (which inhibits BCL-2, BCL-X_L_ and BCL-W) (50), probably because BCL-2 levels are too low. However, BCL-2 levels are high in the highly aggressive subset of human Diffuse Large B-cell lymphomas (C3 DLBCLs) that harbor BCL-2 as well as MYC translocations or overexpress BCL-2 and MYC via other mechanisms (51,52). To model such ‘Double-Expressor’ DLBCLs and test their sensitivity to ABT-199 in the presence and absence of MNT, we infected *Mnt^fl/fl^ Eμ-Myc/CreERT2* lymphoma CLs with our *hBCL-2/GFP* retrovirus (35). Immunophenotyping analysis (data not shown) confirmed that the resulting lines remained B lymphoid, which differentiates them from the lympho-myeloid progenitor cell lymphomas that develop in *Eμ-Myc/Eμ-BCL-2* bi-transgenic mice (14,53).

Western blot analysis showed that these *BCL-2^hi^ Mnt^fl/fl^ Eμ-Myc/CreERT2* CLs had higher BIM and PUMA and, often, lower MCL-1 levels than control GFP-virus infected CLs, and that 4-OHT-mediated *Mnt* deletion resulted in even higher levels of BIM and PUMA (Fig. 6A and Supplementary Fig S6A). Of note, the high BCL-2 and reduced MCL-1 expression rendered these *Eμ-Myc* lymphoma cells dependent on ectopic BCL-2 and they were now resistant to MCL-1 inhibitor (S63845) (data not shown).

**Figure 6.**
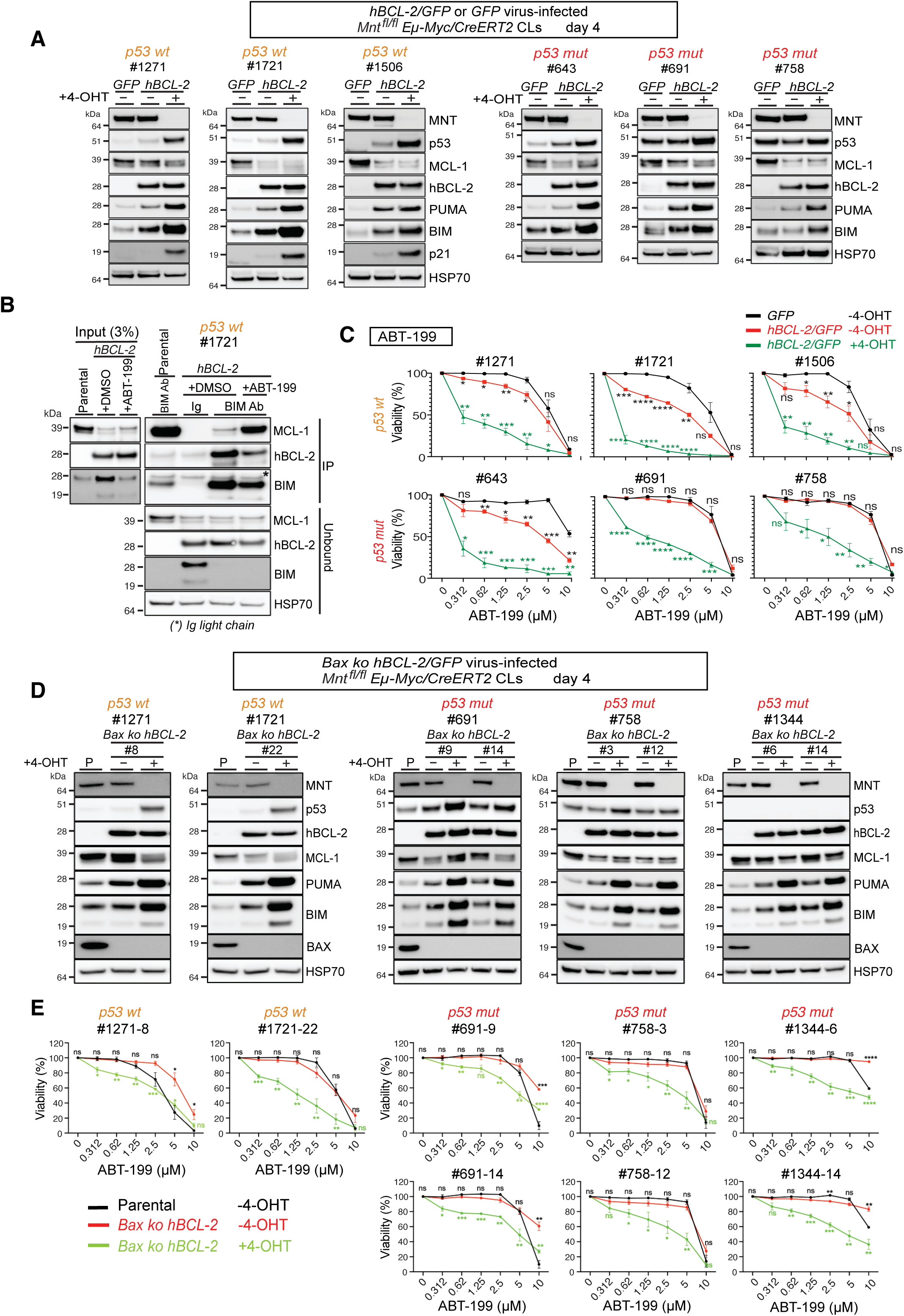
MNT loss increases the sensitivity of *Eμ-Myc* lymphoma cells overexpressing BCL-2 to the BCL-2-specific BH3 mimetic ABT-199 (venetoclax). **A**, Western blot expression analysis of MNT, p53 and indicated BCL-2 protein family members before and after 4-OHT-mediated *Mnt* deletion. The indicated *p53 wt* and *p53 mut Mnt^fl/fl^ Eμ-Myc/CreERT2* lymphoma CLs were infected with *hBCL-2/GFP* or control *GFP* retrovirus and GFP-positive cells selected by FACS to establish sub-lines (see Methods). Lymphoma CLs were incubated in medium +/-4-OHT for 20 hr, then diluted 8-fold and harvested on day 4 for western blot analysis using HSP70 as a loading control. **B,** BCL-2-bound BIM decreases and MCL-1-bound BIM increases in BCL-2 over-expressing *Eμ-Myc/CreERT2* lymphoma cells following treatment with ABT- 199. h*BCL-2* virus-infected (*p53 wt Eμ-Myc* lymphoma CL #1721 was treated with ABT-199 (5 μM) for 12 hr prior to immunoprecipitation of lysates with BIM or control Ig antibody, followed by western blot analysis with antibodies for MCL-1, hBCL-2 and BIM (see Methods). **C,** Sensitivity of parental and BCL-2 over-expressing *Eμ-Myc/CreERT2* lymphoma CLs to ABT-199 before and after *Mnt* deletion. Indicated lymphoma CLs infected with *GFP* or *hBCL-2/GFP* retrovirus (*BCL-2^hi^*) were treated +/- 4-OHT (0.5 μM), followed by dilution, then treated on day 4 with different concentrations of ABT-199 for 24 hr before determining viability by flow cytometry. Black curves show viability of control (4-OHT untreated) *Mnt^fl/fl^ GFP* virus-infected lymphoma CLs, and green and red curves show viability of *Mnt^fl/fl^ BCL-2/GFP* virus-infected lymphoma CLs treated with 4-OHT or medium alone respectively. Black asterisks indicate significance of differences in sensitivity due to overexpression of BCL-2 alone ie red curves vs black curves. Green asterisks indicate significance of differences in sensitivity due to MNT loss for *BCL-2^hi^*lymphoma CLs ie green curves vs red curves. Results are shown for at least 3 independent experiments and plotted as mean ± SEM; *P≤ 0.05, **P≤ 0.01, *** P≤ 0.001, **** P≤ 0.000; ns= not significant. **D, E,** MNT loss enhances ABT-199 sensitivity of *BCL-2^hi^ Eμ-Myc* lymphoma cells even after BAX loss. Independent *Bax ko* clones of *p53 wt* and *p53 mut Eμ-Myc/CreERT2* lymphoma CLs (see Supplementary Fig S6D) were infected with the *hBCL-2/GFP* retrovirus. **D** shows expression of MNT, p53 and the indicated apoptosis regulators in *Bax ko BCL-2^hi^* clones on day 4 following brief treatment +/- 4-OHT (see text) compared to the untreated parental line (P). Blots are typical of 2 independent experiments using different lysates for indicated clone of each lymphoma CL. **E** shows sensitivity of the indicated CLs to 24 hr treatment with ABT-199 over a range of concentrations. Black curves show viability of indicated control (4-OHT untreated) parental lymphoma CLs, and pale green and red curves show viability of *Bax ko BCL-2^hi^ Mnt^fl/fl^ Eμ-Myc* lymphoma CLs treated with 4-OHT or medium alone respectively. Black asterisks indicate significance of differences in sensitivity between parental and 4-OHT untreated *Bax ko BCL2^hi^ Eμ-Myc/CreERT2* cells. Green asterisks indicate significance of differences in sensitivity due to MNT loss for *Bax ko BCL-2^hi^ Eμ-Myc/CreERT2* lymphoma clones ie green curves vs red curves. Results are shown for at least 3 independent experiments and plotted as mean ± SEM; *P≤ 0.05, **P≤ 0.01, *** P≤ 0.001, **** P≤ 0.000; ns= not significant.

BIM immunoprecipitation performed with *p53 wt* and *p53* mutant CLs (Fig. 6B; Supplementary S6C) revealed that most BIM was bound to BCL-2 in the *BCL-2^hi^*CLs rather than to MCL-1 as in the parental CLs (eg compare lanes 4 and 6 in Fig. 6B). The reduced level of MCL-1 in many *BCL-2^hi^* CLs likely reflects lower stability of MCL-1 unbound to BIM or other BH3-only proteins. However, after 12-hr treatment with the BCL-2-specific BH3 mimetic ABT-199 (venetoclax), the level of BCL-2 bound to BIM decreased, indicative of preferential binding of ABT-199 to BCL-2, whereas the level of MCL-1 complexed with BIM increased (compare lanes 6 and 7 in **Fig. 6A**, and lane 7 with lanes 8 and 9 in **Fig. S6C**), as reported earlier (54).

After *Mnt* deletion, the apoptosis-resistant *BCL-2^hi^ Eμ-Myc/CreERT2* lymphoma CLs proliferated more slowly than their parental counterparts (Supplementary Fig. S6B), as shown above for *MCL-1^hi^* or *BaxBak dko Mnt^fl/fl^ Eμ-Myc/CreERT2* lymphoma cells.

Next, we tested the response of the *BCL-2^hi^ Eμ-Myc/CreERT2* lymphoma CLs to ABT-199 before and after *Mnt* deletion. Cells treated briefly with 4-OHT on d1 were incubated on d4 with increasing concentrations of ABT-199 for 24 hr, as described in Fig. S4A. As expected from their low level of endogenous BCL-2, all control *GFP-*virus-infected *Eμ-Myc/CreERT2* CLs were very resistant to ABT-199 (Fig. 6C, black curves). BCL-2 over-expression rendered 3 of 3 *p53* wt CLs sensitive, but only 1 of 3 *p53* mutant lines (compare red to black curves). Importantly, however, MNT loss dramatically increased the sensitivity of all *p53* wt and all *p53* mutant lines to ABT-199 (compare green with red curves), presumably due to the further upregulation of BIM and PUMA (Fig. 6A).

### Impact of MNT loss in ‘Double Expressor’ DLBCLs lacking BAX

Loss of BAX function confers resistance to venetoclax treatment in B cell lymphomas (45,55,56) and acute myeloid leukemia (AML) (57). To determine if MNT loss could enhance the venetoclax sensitivity of BAX-deficient ‘Double Expressor’ DLBCLs, we first generated *Bax ko* clones of several *Mnt^fl/fl^ Eμ-Myc/CreERT2* lymphoma CLs, using CRISPR/Cas9 (Supplementary Fig. S6D). Both *p53* wt and *p53* mutant *Bax ko* clones remained sensitive to apoptosis induced by MNT loss (Supplementary Fig. S6D), indicating that BAK is sufficient to mediate apoptosis triggered by MNT loss. Next, we infected these *Bax ko Mnt^fl/fl^ Eμ-Myc/CreERT2* clones with the *hBCL2/GFP* virus and determined the expression of apoptosis regulators (Fig. 6D). Of note, there was a major increase in the level of PUMA and BIM 4 days after 4-OHT-induced *Mnt* deletion (compare – and + tracks).

Like their parental lines (black curves), the cloned *Bax ko BCL-2^hi^* CLs were very resistant to ABT-199 (red curves) (Fig. 6E). Notably, however, MNT loss significantly improved their sensitivity (compare green to red curves), presumably at least in part due to the higher levels of PUMA and BIM after *Mnt* deletion.

Taken together, these data suggest that targeting MNT function would be an effective strategy for improving venetoclax treatment of ‘Double Expressor’ DLBCLs, even those that have evolved to be resistant by losing p53 and/or BAX function.

### Impact of *MNT* Deletion in Burkitt Lymphoma CLs

To extend our studies to MYC-driven human lymphomas, we investigated CLs established from human Burkitt lymphomas. Fig. 7A compares the expression of apoptosis regulators in 3 long-established lines: BL-2, which is wt for *p53* but carries a homozygous deletion of *p14^ARF^* (58), and two *p53* mutant CLs, Rael-BL and BL-41. All three lines have robust levels of anti-apoptotic MCL-1 and BCL-X_L_ but only BL-41 also has strong BCL-2 expression. All express pro-apoptotic BAX and BAK but the major pro-apoptotic BH3-only protein BIM is undetectable in BL-2, as reported previously (59).

**Figure 7.**
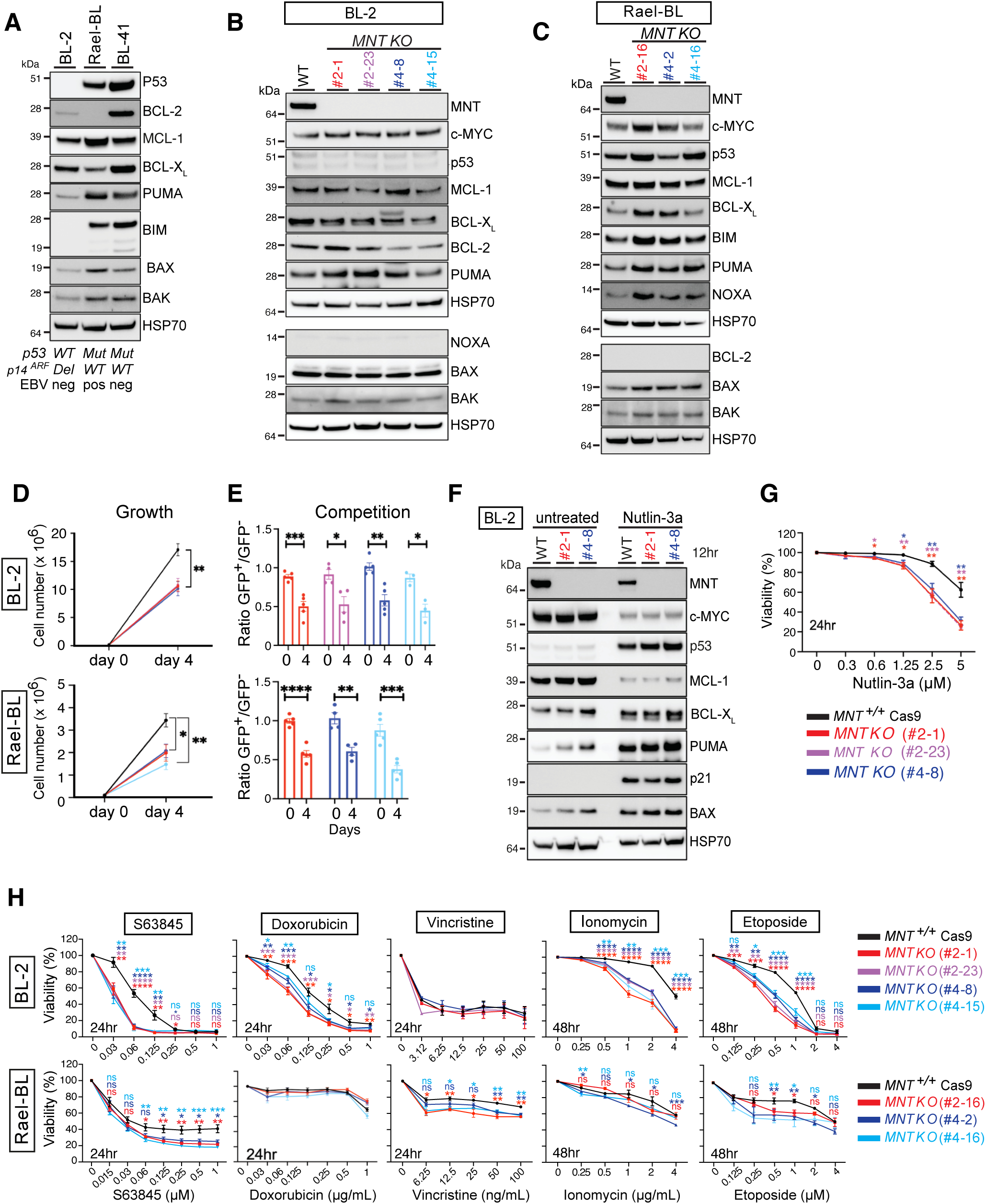
Impact of MNT loss in Burkitt Lymphoma cell lines. **A**, Western blot expression analysis of indicated apoptosis regulators in Burkitt Lymphoma cell lines BL-2, Rael-BL and BL-4. HSP70 was used as a loading control. Published genotypes for *p53*, *p14^ARF^* and Epstein Barr Virus (EBV) are indicated below western. **B,** Western blot expression analysis of the indicated apoptosis regulators in parental BL-2 cells and independent clonal *MNT KO* derivatives. Clones #2.1 and # 2.23 were derived by CRISPR/Cas9 (see Methods) using sgRNA #2, and clones #4.8 and #4-15 were derived using sgRNA #4 (see Supplementary Fig.S7A). HSP70 was used as a loading control. **C**, Western blot expression analysis of the indicated apoptosis regulators in parental Rael-BL cells and independent clonal *MNT KO* derivative lines. Clone#2.16 was derived by CRISPR/Cas9 using sgRNA #2 and clones #4.2 and #4-16 were derived using sgRNA #4. **D**, MNT loss reduces cell proliferation in BL-2 and Rael-BL cells. 10^5^ cells were cultured in 5mL and counted on day 4. Note the slower growth of the Cas9^+^ parental Rael-BL compared to Cas9^+^ parental BL-2 cells (see y axis). Black lines indicate Cas9^+^ parental cell lines and coloured lines indicate derivative *MNT KO* clones (see key). Results are shown as mean of three independent experiments ± SEM *P≤ 0.05, **P≤ 0.01. **E**, Cell competition assay confirming that *MNT KO* clones of BL-2 and Rael-BL grow more slowly than their Cas9^+^ parental counterparts. 10^5^ viable cells of *MNT KO* (mCherry^+^GFP^+^) and parental (mCherry^+^GFP^-^) BL-2 (upper panel) and Rael-BL (lower panel) CLs were mixed together and cultured in 5 mL for 4 days. The relative proportion of viable GFP^+^ and GFP^-^ cells in the cultures was determined on day 0 and day 4 by flow cytometry. Results are shown as mean of three independent_experiments (dots) +/- SEM; *P≤ 0.05, **P≤ 0.01, *** P≤ 0.001, **** P≤ 0.000. **F** MDM-2 inhibitor Nutlin-3a stabilizes wt p53 protein in BL-2 cells. Cas9^+^ parental and two indicated *MNT KO* BL-2 cell lines were incubated for 12 hr +/- Nutlin-3a (5 μM) in the presence of Q-VD-OPh (to prevent caspase activation and cell demolition). Western blot analysis shows that Nutlin-3a treatment increased p53 protein, accompanied by increased expression of its direct targets PUMA, p21 and BAX. In addition, BCL-X_L_ was increased, but MCL-1 and MYC were decreased. HSP70 was used as a loading control. Blot is typical of 3 independent experiments. **G**, *MNT KO* BL-2 clones show lower viability than the Cas9^+^ parental BL-2 cells after stabilization of p53 by Nutlin-3a. Cells were treated with Nutlin-3a (5 μM) for 24 hr, stained with DAPI and cell viability was determined by flow cytometry. **H**, Comparison of sensitivity of Cas9^+^ parental and *MNT KO* BL-2 and Rael-BL CLs to the MCL-1 inhibitor S63845, Doxorubicin, Vincristine, Etoposide and Ionomycin. Cells were treated for 24 or 48 hr with the indicated agents at increasing concentrations, then stained with DAPI to determine cell viability by flow cytometry. Three independent experiments were performed and the results are presented as mean ± SEM. Asterisks color-coded for the different lines (see key) indicate significant differences in sensitivity at different concentrations; *P≤ 0.05, **P≤ 0.01, *** P≤ 0.001, **** P≤ 0.0001, ns not significant.

To delete MNT from these lines, we utilised a dual lentiviral vector system (60) to introduce *Cas9* and two independent sgRNAs located in exon 2 (Supplementary Fig. S7A). When treated with doxycycline to induce sgRNA expression, only BL-41 cells showed reduced viability on d4, accompanied by reduced MNT (Supplementary Fig. S7B). We attempted to isolate *MNT KO* clones from doxycycline-treated cultures, succeeding with BL-2 and Rael-BL but not with BL-41, perhaps because *MNT*-deleted BL-41 cells died too fast. Western blot analysis confirmed the absence of full-length MNT protein in three Rael-BL and four BL-2 clones (Fig. 7B, C), and although three of the BL-2 clones expressed small N-terminal polypeptides of 19 to 29 kD, presumably N-terminal readthrough products, these were confined to the cytoplasm and would therefore be expected to be non-functional (Supplementary Fig. S7C, D).

We assume that the *MNT KO* BL-2 and Rael-BL clones carry mutations that enabled them to resist apoptosis triggered by MNT loss. In BL-2, survival capacity would have been strongly enhanced by the pre-existing mutations inactivating *BIM* and *p14^ARF^*(Fig. 7A). No other consistent differences in apoptosis regulators were detected in the *MNT*-deleted clonal derivatives of BL-2 (Fig. 7B). The basis for the apoptosis-resistance of *MNT KO* Rael-BL cells in the face of their increased expression of BIM, PUMA and NOXA proteins (Fig. 7C) and mRNA (Supplementary Fig. S7E) is unclear.

Despite their unchanged level of c-MYC (Fig. 7B), the *MNT KO* clones of both BL-2 and Rael-BL grew more slowly than their Cas9+ parental cells (Fig. 7D) and this was confirmed by competition assays between the *MNT KO* (GFP^+^) cells with the Cas9^+^ parental (GFP^-^) cells (Fig 7E and Supplementary S7F). As a negative control, GFP^+^ *hMNT-sgRNA* clones that had not been treated with doxycycline grew at the same rate as parental cells (Supplementary Fig. S7G,H).

To test the functionality of p53 in BL-2, we used Nutlin-3a, which stabilises p53 by inhibiting its interaction with the E3 ligase MDM2 (33). As expected, Nutlin3a induced high levels of p53 protein and its targets PUMA, p21 and BAX in both Cas9^+^ parental and *MNT KO* BL-2 cells (Fig. 7F). Importantly, the *MNT KO* BL-2 clones were more susceptible to p53-mediated apoptosis induced by Nutlin-3a than the Cas9+ parental line (Fig. 7G), suggesting that MNT loss would enhance sensitivity to anti-cancer drugs that induce p53.

When the drug sensitivity of the Cas9^+^ parental cells (black curves) and independent *MNT KO* clones were compared (Fig. 7H), we found that MNT loss significantly enhanced the sensitivity of BL-2 cells to killing by the DNA-damaging drugs Doxorubicin, and Etoposide, and also to Ionomycin, but had little impact on their response to Vincristine, to which they were already highly sensitive. Although the (EBV-positive, *p53* mutant) Rael-BL cells were more drug-resistant than BL-2, MNT loss significantly improved their responsiveness to Vincristine and Etoposide. For both BL-2 and Rael-BL, MNT loss also significantly increased sensitivity to the MCL-1 inhibitor S63845.

## DISCUSSION

The central importance of the MYC transcription factor in cancer development is well accepted. However, the complexity of the interactions that occur between members of the wider MYC family that determine transcriptional and phenotypic outcomes in different normal and neoplastic cell types is only just beginning to be appreciated eg (61). Amongst the MYC-related repressors, MNT is the most highly conserved and widely expressed. A mouse model of MYC-driven breast cancer suggested that MNT was a tumor suppressor (23). However, the Hurlin lab and ours have shown that, in lymphomagenesis, MNT synergises with MYC, by repressing apoptosis (24–27). We have also recently shown that AML driven by *MLL::AF9* is MNT-dependent (62).

In this study, using CLs derived from multiple independent *Mnt*-deletable *Eμ-Myc* lymphomas and human Burkitt lymphomas, we have tested the MNT dependency of fully malignant MYC-driven cells, further explored MNT’s physiological roles and provided preclinical proof of concept that MNT inhibitors would facilitate lymphoma treatment.

We showed that both p53 wt and p53 mutant *Eμ-Myc* lymphoma cells undergo apoptosis following MNT loss and have extended understanding of the mechanism. We previously found that MNT loss in premalignant lymphoid cells elevated the level of the pro-apoptotic BH3-only protein BIM (26,27). Here we have shown that MNT loss also elevates p53 and the BH3-only protein PUMA (we lacked a suitable antibody to test for NOXA), and that PUMA is increased by both p53-dependent and p53-independent mechanisms. Furthermore, our analysis of CUT&RUN data from B220^+^ splenic B cells published by the Eisenman lab (37) suggests that *Bim*, *Puma* and *p53* genes are all direct targets of MNT/MAX heterodimers. The increased BIM, PUMA and NOXA proteins found in *MNT KO p53* mutant Rael-BL cells implies that MNT also suppresses expression of BH3-only proteins in human lymphoid cells.

By analysis of *Eμ-Myc* lymphoma cells engineered to be apoptosis-resistant, we also showed that MNT loss inhibited cell proliferation and promoted senescence. Our results suggested that MNT loss mediates senescence programs by both a p53-dependent pathway (via p21) and a p53-independent pathway (likely p16). Like *Mnt-*deleted *Eμ-Myc* lymphoma cells, *MNT KO* clones of BL-2 and Rael-BL proliferated more slowly than their parental cells regardless of *p53* status and/or other genetic differences such as EBV status. These observations add further weight to our argument for MNT as a therapeutic target (26,27), since lymphoma cells that acquire mutations enabling them to escape apoptosis after MNT inhibition would nevertheless have poor expansion potential due to their limited proliferative capacity.

To provide preclinical proof-of-concept for developing drugs that inhibit MNT function, we treated our panel of *Mnt*-deletable *Eμ-Myc* lymphoma CLs with diverse cancer drugs, including the MCL-1 specific BH3 mimetic S63845, as *Eμ-Myc* lymphomas are known to be MCL-1 dependent (34). The responsiveness of these CLs to the drugs would be expected to vary, depending on the nature of acquired synergistic oncogenic mutations (largely unidentified) and whether they are refractory to apoptosis by virtue of being *p53* mutant or engineered to over-express hMCL-1. We found that inducing MNT loss prior to drug treatment significantly enhanced the sensitivity of three of five *p53* mutant *Mcl-1^wt^ Eμ-Myc* lymphoma CLs to S6385, Cisplatin and Doxorubicin but not Vincristine, to which they were already very sensitive. MNT loss also sensitised three of three *p53* wt *hMCL-1^hi^*CLs to Cisplatin, Doxorubicin and Vincristine, as well as to S63845. Of the doubly apoptosis-refractory *hMCL-1^hi^ p53* mutant CLs, MNT loss increased the sensitivity of 2 of 4 lines tested to Cisplatin, 4 of 4 to Vincristine and 3 of 3 to S63845 but none of four to Doxorubicin. Finally, S63845-resistant *p53* mutant sub-lines, generated by culturing the cells in increasing levels of this drug, became more sensitive to Cisplatin, Doxorubicin and Vincristine after MNT loss. We surmise that a major reason for the increased sensitivity to all these anti-cancer agents is the increase in the level of BH3-only proteins such as BIM and PUMA after MNT loss.

Because they express little BCL-2, *Eμ-Myc* lymphoma cells are highly resistant to the BCL-2-specific BH3-mimetic ABT-199 (venetoclax). However, when engineered to express high level of BCL-2, we found that *Mnt* deletion rendered all the CLs highly sensitive to ABT-199, regardless of their p53 status, even after loss of BAX, a mutation observed in patients that become venetoclax-resistant (45,55,56). We ascribe the improved venetoclax sensitivity to the increase in pro-apoptotic BIM and PUMA found in these lymphoma cells after *Mnt* deletion. These results suggest that a MNT inhibitor would greatly improve venetoclax therapy for the highly aggressive human ‘double-expressor’ DLBCLs (51).

To extend our findings to MYC-driven human lymphoma cells, we also studied the impact of MNT deletion in three Burkitt lymphoma CLs, using CRISPR/Cas-9. The long culture history of these CLs has presumably facilitated the acquisition and selection for multiple alterations that enhance cell survival. Doxycycline-induced expression of sgRNAs targeting *MNT* resulted in reduced cell viability for BL-41 cells but not BL-2 or Rael-BL. Of note, *MNT KO* BL-2 and Rael-BL cells were more sensitive to the MCL-1 inhibitor S63845 than their parental cells. Furthermore, in BL-2, despite the absence of BIM (59), MNT loss rendered the cells more sensitive to doxorubicin, etoposide and ionomycin. *MNT KO* Rael-BL cells were more sensitive to vincristine and etoposide.

Taken together, our results present a strong case for MNT as an attractive target for non-Hodgkin’s lymphomas and other MYC-driven blood cancers. Rather than trying to inhibit MYC, we propose a strategy that takes advantage of MYC’s innate capacity to induce apoptosis and amplifies that drive, by preventing MNT from suppressing genes that promote apoptosis in tumor cells expressing high MYC levels. Our preclinical data indicate that MNT inhibitors would promote apoptosis in treatment-naïve MYC-driven lymphomas and also increase their sensitivity to both conventional chemotherapeutic agents and targeted therapies such as BH3 mimetic drugs.

While up-regulation of pro-apoptotic proteins is undoubtedly a major factor for the increase in apoptosis in the absence of MNT, other pathways may play an important role. MNT can compete not only with MYC/MAX heterodimers that bind to E-box sites to regulate cell proliferation, it also competes MLX heterodimers that bind ChoRE sites and regulate genes controlling metabolism (19,22). Thus, MNT loss also might alter metabolic pathways that render lymphoma cells more vulnerable to apoptosis.

## METHODS

### Mice

Mice used here were *Eμ-Myc* (3), *Rosa26CreERT2* (32) (hereafter *CreERT2*) and *Mnt^fl/+^* (63) (kind gift of PJ Hurlin in 2005). All were on a C57BL/6 background, bred and monitored at the Walter and Eliza Hall Institute (WEHI) under the supervision of trained veterinarians and treated and analyzed in accordance with WEHI’s animal ethics committee regulations.

### Eμ−Myc/CreERT2 and Burkitt Lymphoma CLs

Cell suspensions prepared from pre-B/B lymphomas arising in *Mnt^fl/fl^*and *Mnt^+/+^ Eμ-Myc/CreERT2* mice were cultured in complete Opti-MEM containing 10% heat-inactivated fetal bovine serum (FBS), 50 μM 2-mercaptoethanol, 20 mM Hepes pH 7.5, 2 mM L-glutamine, 1 mM sodium pyruvate, 1X MEM non-essential amino acids, 100 U/mL penicillin and 100 U/mL streptomycin, changing the medium every 3 days until the CL was established and cryopreserved. Subsequently, the lymphoma CLs were cultured in complete high glucose Dulbecco’s Modified Eagles’ medium (DMEM) containing (10% FBS, 50 μM 2-mercaptoethanol, 20 mM Hepes, 2 mM L-glutamine, 1 mM sodium pyruvate, 1X MEM non-essential amino acids, 100 U/mL penicillin and 100 U/mL streptomycin).

Human Burkitt’s Lymphoma CLs (BL-2, Rael-BL and BL-41), authenticated by short-tandem-repeat (STR) profiling at the Australian Genome Research Facility, were maintained in RPMI-1640 medium containing 10% FBS, 50 μM 2-mercaptoethanol, 2 mM L-glutamine, 1 mM sodium pyruvate, 1X MEM non-essential amino acids, 100 U/mL penicillin and 100 U/mL streptomycin.

All lymphoma CLs were incubated at 37°C with 5% CO_2_ and maintained in culture for a maximum of 8 weeks before accessing a new frozen vial. Regular mycoplasma screening was conducted using Lonza’s MycoAlert kit (Cat #LT07-118).

2-mercaptoethanol was purchased from Sigma-Aldrich (Cat #M3148) and all other reagents from Thermo Fisher: Opti-MEM (Cat #31985070), DMEM (Cat #11965092), RPMI-1640 (Cat #11875093), Hepes 1 M pH 7.5 (Cat #15630080), L-glutamine 200 mM (Cat #25030081), sodium pyruvate 100 mM (Cat #11360070), MEM non-essential amino acids solution 100X (Cat #11140050) and penicillin-streptomycin 10,000 U/mL (Cat #15140122).

### Nutlin-3a Treatment

To determine the *p53* status of *Eμ-Myc/CreERT2* lymphoma CLs, cells were treated with 5 μM Nutlin-3a (Cayman Chemical #18585) for 24 hr and viability was determined by flow cytometry (see below). Nutlin-3a inhibits MDM2, stabilising wt p53(33). *p53* wt lymphoma cells undergo apoptosis in response to Nutlin-3a but *p53* mutant or *p53* knockout cells are resistant(64).

To determine the p53 response in BL-2 cells, 2x10^6^ Cas9^+^ parental BL-2 and *MNT-KO* BL-2 clones were incubated with Nutlin-3a (5 μM) or DMSO for 12 hr in presence of the caspase inhibitor Q-VD-OPh (ChemSupply Australia, Cat #GK2336). Lysates of treated cells were analyzed by western blot.

### Generation of Eμ-Myc/CreERT2 Lymphoma CLs Expressing hMCL-1 or hBCL-2

Retroviral vectors expressing human BCL-2 or FLAG-tagged human MCL-1 (35), constructed in our lab by Dr Cassandra Vandenberg, were used to derive apoptosis-resistant *Eμ-Myc/CreERT2* lymphoma CLs. To prepare retrovirus-containing supernatants, HEK293T cells (0.5 x 10^6^) were seeded in 3 mL DMEM containing 10% FBS in 6-well plates, incubated at 37° C for 24 hr, then transfected with 4 μg of empty vector *pMIG GFP* (Addgene, plasmid #9044), *pMIG-Flag-hMCL-1/GFP* or *pMIG-hBCL2/GFP* DNA, together with 2 μg retroviral *GAG/POL* packaging plasmid (Addgene, Plasmid #14887) and 1 μg ENV packaging plasmid (Addgene, Plasmid # 8454), using the calcium phosphate protocol. After incubation for 24 hr, the medium was replaced and the virus-containing supernatant was collected 48 hr later. For spinfection, cells (0.5x10^6^ in 1 mL) were mixed with retrovirus-containing supernatant (3 mL) in polybrene (4 μg/mL), centrifuged at 2500 rpm for 45 min at 32°C and then incubated overnight at 37°C. The medium was replaced 24 hr after infection and, after another 2 days, GFP-positive cells were selected using a FACS Cell sorter Aria^TM^ III (BD). The same protocol was used to generate *Bax ko Mnt^fl/fl^ Eμ-Myc/CreERT2* clones expressing hBCL2.

### Generation of Mnt^fl/fl^ Eμ-Myc/CreERT2 Lymphoma CLs Resistant to MCL-1 Inhibitor

*Mnt^fl/fl^ Eμ-Myc/CreERT2* lymphoma cells were cultured in increasing concentrations of S63845 (0.35 μM to 0.5 μM to 1 μM to 2 μM). At each step, lymphoma cells were cultured until >70% cells were viable before the higher drug dose was added. Resistance to S63845 was confirmed using cell survival assays at 1 μM S63845.

### 4-OHT-Mediated Mnt Deletion in Mnt^fl/fl^ Eμ-Myc/CreERT2 Lymphoma CLs in vitro

Cells were cultured at 37°C for 20 hr in complete DMEM medium (2x10^5^ cells in 1 mL) containing 4-OHT (0.5 μM) (Sigma, Cat #H7904) or medium alone. After 20 hr, 7 mL fresh medium was added to the culture to dilute 4-OHT to 0.06 μM. Viability, PCR and Western blotting analyses were performed at 24, 48, 36, 96 hr and, after further culture of 2x10^5^ viable cells in fresh DMEM, on d8. Cells were stained with Trypan Blue (Invitrogen Cat# T10282) and viable cells (Trypan Blue-negative) counted using a TC20^TM^ Automated Cell Counter (BioRad).

### Tamoxifen-Mediated Mnt-Deletion in Transplanted Mnt^fl/fl^ Eμ-Myc/CreERT2 Lymphoma Cells

Tamoxifen stock (60 mg/mL) (Sigma, Cat #T5648) was prepared in peanut oil containing 10% ethanol. Each lymphoma CL (Ly5.2^+^) was injected iv into six non-irradiated C57BL/6-Ly5.1^+^ recipient mice (10^6^ cells/mouse) and, 5 days later, three recipients were treated by oral gavage on three successive days with tamoxifen (200 mg/kg body weight) and the other three were treated with vehicle. Recipient mice were monitored and euthanized when showing signs of sickness (e.g. hunched stance, increased respiration or enlarged lymphoid organs). Ethical endpoint was determined by experienced animal technicians who were unbiased to the outcome of the experiments. Peripheral blood was analyzed using an ADVIA hematology analyzer (Bayer). Enlarged lymphoid organs (thymus, spleen or LNs) were collected and cryopreserved in FBS containing 10% DMSO. Donor-derived (Ly5.2^+^) CD19^+^ lymphoma cells from enlarged lymphoid organs were purified by FACS sorting and analyzed by PCR genotyping and Western blotting for *Mnt* gene deletion and loss of the MNT protein, respectively.

### PCR Genotype Analysis of 4-OH-T Treated Eμ-Myc/CreERT2 Lymphoma CLs

Cells were lysed in 500 μL lysis buffer (10 mM Tris-HCl pH 8.0, 10 mM EDTA pH 8.0, 0.5% SDS) containing proteinase-K (dilution 1:1000, Sigma-Aldrich, Cat #P4850) and incubated at 56° C overnight. Genomic DNA was precipitated by addition of 500 μL isopropanol and the pellet was washed twice with 70% EtOH before resuspension in TE buffer (10mM Tris-HCl pH 8.0, 1 mM EDTA). DNA concentration was determined using a spectrophotometer (DeNovix DS11). Then 20 ng DNA was added to 19 μL GoTaq Green Master Mix (Promega, Cat #M7123) containing the oligonucleotide primers (final concentration 0.25 μM). The PCR program was 94° C for 3 min followed by 30 cycles (94° C for 30 sec, 58° C for 30 sec, 72° C for 40 sec) and finally 72° C for 5 min. PCR products were separated by gel electrophoresis on a 2% DNA grade agarose gel (Bioline, Cat #BIO-41025) in TAE buffer (40 mM Tris acetate, 1 mM EDTA pH 8.0) containing ethidium bromide (Sigma-Aldrich Cat # 111608) (0.2 μg/mL final concentration) and imaged using a Gel DOC^TM^XR+Gel Documentation system (Bio-Rad). The same protocol was used to identify *Mnt* deletion of *Mnt ko* clones isolated on d8 after treatment with 4-OHT. Primers used and their product size were:

***Eμ-Myc:*** ∼900 bp

*Myctg1*: 5’-CAGCTGGCGTAATAGCGAAGAG-3’
*Myctg2*: 5’-CTGTGACTGGTGAGTACTCAACC-3’

***FloxMnt***: *wt* allele 178 bp, *flox* allele 579 bp

*Mnt CKO-2*: 5’-GTCTCAAGTCGTGGGCATTG-3’
*Hygro R-S*: 5’-GATGTAGGAGGGCGTGGATA-3’
*Hygro R-R*: 5’-GATGTTGGCGACCTCGTATT-3’
*Mnt Seq1*: 5’-CAGATTCAGTGTCCCCTGCT-3’

***DelMnt***: ko 386 bp

*Mnt ko3*: 5’-CAGGTCCTCCAAAAGAGCAG-3’
*Mnt ko4*: 5’-GGAGCAATGTGGAGAGAAGC-3’

***RosaCreERT2***: *Rosa* wt allele 700 bp, *RosaCreERT2* allele 450 bp

*RosaS se*: 5’-GCCAATGCTCTGTCTAGGGGTTGG-3’
*RosaS as*: 5’-CTTGCTCTCCCAAAGTCGCTCTGAG-3’
*RosaS Cre as*: 5’-TCGTTGCATCGACCGGTAATGCAGGC-3’

### CRISPR/Cas9 Gene Editing of Burkitt Lymphoma and Eμ-Myc/CreERT2 Lymphoma CLs

Gene editing was performed using the dual lentiviral vector system developed by Aubrey et al (60). Two independent sgRNAs targeting human *MNT* were cloned into the GFP-expressing doxycycline-inducible FgH1tUTG vector (Addgene, plasmid #70183). To produce lentivirus supernatant, HEK 293T cells were co-transfected with packaging vectors (1 μg pCMV-Rev, 1.5 μg pMDLg/pRRE and 1 μg pCMV-VSV-G) and with 1.5 μg Cas9-mCherry vector (pFUGW-Cas9-mCherry) (Addgene, plasmid #70182)(60) or with 1.5 μg sgRNA vector using the calcium phosphate protocol. After incubation for 24 hr, the medium was replaced and lentivirus containing supernatants were harvested 2 days later.

To derive *MNT KO* BL-2 and Rael-BL CLs, cells were infected with the Cas9 lentivirus and FACS sorted for mCherry^hi^ cells, which were then infected with the sgRNA lentivirus, using spinfection as above. To induce sgRNA expression, mCherry^+^ GFP^+^ cells were treated with 1 μg/mL doxycycline (Sigma, Cat #D9891), incubated at 37°C for 96 hr and FACS sorted to isolate single cells. Western blotting was used to identify *MNT KO* clones.

To derive *Bax ko* and *BaxBak dko Eμ-Myc/CreERT2* CLs, Dr Sarah Diepstraten (WEHI) kindly provided mouse *Bax* and *Bak* sgRNAs cloned in the *U6-gRNA/PGK-Puro-2A-BFP* lentivirus vector, obtained from the Sanger Arrayed Whole Genome Lentiviral CRISPR Library (Merck, Cat #MSANGRG). Production of lentivirus and infection were as described above. After infection with Cas9 lentivirus, sorted mCherry^hi^ *Eμ*-*Myc/CreERT2* lymphoma cells were incubated for 24 hr with supernatants containing sgRNA lentivirus and immediately subjected to limited dilution to isolate single cells (mCherry^+^ BFP^+^). Western blotting was used to identify *Bax ko* or *BaxBak dko* clones.

sgRNA sequences used were:

mouse *Bax*: 5’- ACAGGGGCCTTTTTGCTACAGG-3’
mouse *Bak:* 5’- GGGCAAGTTGTCCATCTCGGGG-3’
human *MNT*: sgRNA#2: 5’-TCCCAGGGGTGGCGCCTCCATGCG-3’
human *MNT*: sgRNA#4: 5’-TCCCCTCCTTAATGCTGAGTCCGG-3’

### cDNA Synthesis and Taq-Man PCR Analysis

Total RNA was purified from 1-2 x 10^6^ viable *Eμ-Myc/CreERT2* lymphoma cells collected on d4 after 4-OHT treatment or from *MNT-wt* and *MNT-KO* human Rael-BL cells, using RNeasy Kits (Qiagen, Cat #74104). RNA was treated with RNase-free DNase set (Qiagen, Cat #79254). cDNA was synthesized using Oligo (dT)12-18 primer (Invitrogen, Cat #18418012) and M-MLV reverse transcriptase (Invitrogen, Cat #28025013) as described by the manufacturer. Real-time PCR was conducted with Taq-man Fast Advanced Master Mix (ThermoFisher, Cat #4444556) using following specific probes: mouse *Bim/Bcl2l11* (Mm00437796_m1) (ThermoFisher, Cat #4331182), mouse *p53* (Mm01731287_m1) (ThermoFisher, Cat #4331182), mouse *Puma/Bbc3* (Mm00519268_m1) (ThermoFisher, Cat #4331182), mouse *p21* (Mm00432448_m1) (ThermoFisher, Cat #4453320), human *BIM (Bcl2l11)* (Hs00708019_s1 (Thermo Fisher, Cat #4331182), human PUMA/BBC3 (Hs00248075_m1) (Thermo Fisher, Cat #4331182), human *NOXA/PMAIP1* (Hs00560402_m1) (Thermo Fisher, Cat #4453320) and, as a loading control, mouse *Gapdh* (Mm999999_g1) (ThermoFisher, Cat #4331182) and human *GAPDH* (Hs02786624_g1) (Thermo Fisher, Cat #4331182). PCR reactions were run on ViiA7 Real-time PCR system (ThermoFisher). For *Eμ-Myc* CLs, the fold change (ΔΔCt) of target gene expression of treated condition (+4-OHT) was normalized to *Gapdh* expression and to carrier condition (-4-OHT) using the 2^-(ΔΔCt)^ method (see below). For Burkitt CLs, the fold change (ΔΔCt) of target gene expression of *MNT-KO* clones was normalized to *GAPDH* expression and to the corresponding Cas9^+^ parental CL. All data are presented as mean ± SEM of at least 3 independent experiments.

ΔCt= Ct (*target gene*) - Ct (*Gapdh*)

ΔΔCt=ΔCt (+4-OHT) - ΔCt (-4-OHT) for *Eμ-Myc* CLs

ΔΔCt=ΔCt (*MNT KO*) - ΔCt (parental CL) for Burkitt CLs

### Apoptosis Assays

4-OHT-treated *Eμ-Myc/CreERT2* cells were centrifuged, washed with Phosphate-Buffered Saline (PBS 1X Thermo Fisher) and stained in Annexin-V binding buffer (BioLegend Cat #422201) containing Annexin-V-Alexa Fluor 647 (made in house, dilution 1:1000) and 1 μg/mL propidium iodide (PI) (Sigma-Aldrich, Cat #P4710) (e.g. Fig S1A); viable cells are PI^-^Annexin-V^-^. Alternatively (e.g. after drug treatment) cell viability was determined by flow cytometry after staining with PI (1 μg/mL) or 4’,6-diamidino-2-phenylindole (DAPI) (5μg/mL); viable cells are PI^-^ or DAPI^-^.

For 4-OHT-treated *BaxBak dko Mnt^fl/fl^ Eμ-Myc/CreERT2* or *Bax ko hBCL-2^hi^ Eμ-Myc/CreERT2* lymphoma cells, which are BFP-positive, cells were stained with PI and analyzed in a Fortessa flow cytometer using the following settings: Blue laser 488 nm – Detector B695/40 detects PI signal, Blue laser 488 nm – Detector B530/30 detects GFP signal and Violet laser 405 nm – Detector V450/50 detects BFP signal. The yellow laser 546 nm was switched off to avoid leakiness of the mCherry signal into the PI signal. See Fig. S6E for representative FACS plots.

### Western Blot Analysis

Western blots were prepared and analyzed as previously (26). Membranes were probed using the antibodies listed below.

| **Antibodies** | **Source** | **Identifier** | **Dilution** |
| --- | --- | --- | --- |
| MNT | Bethyl Laboratories | #A303-626A | 1/2000 |
| MAX | Abcam | #ab199489 | 1/2000 |
| c-MYC | Cell Signaling | #5605 clone D84C12 | 1/2000 |
| mouse p21 | Cell Signaling | #64016 | 1/1500 |
| human p21 | Cell Signaling | #2947 clone 12D1 | 1/1500 |
| mouse p19-Arf | Rockland | #200-501-891 | 1/1500 |
| mouse p16-INK4A | Cell Signaling | #29271 clone E5F3Y | 1/2000 |
| mouse p53 | Novocastra Leica Biosystems | #NCL-Lp53-CM5p clone CM5 | 1/2000 |
| human p53 | Santa Cruz | #sc-126 | 1/2000 |
| Suv39h1 | Cell Signaling | #8729 clone D11B6 | 1/1500 |
| MCL-1 | WEHI mAb Lab | clone 19C4-15 | 1/2000 |
| mouse BCL-2 | WEHI mAb Lab | clone hamster 3F11 | 1/2000 |
| human BCL-2 | WEHI mAb Lab | clone Bcl-2-100 | 1/2000 |
| BCL-X<sub>L</sub> | Cell Signaling | #2764 clone 54H6 | 1/2000 |
| BIM | Cell Signaling | #2933 clone C34C5 | 1/2000 |
| PUMA | Cell Signaling | #986782 clone E2P7G | 1/2000 |
| human NOXA | Cell Signaling | #14766 clone D8L7U | 1/1000 |
| BAX | Cell Signaling | #2772 | 1/1000 |
| BAK | WEHI mAb Lab | clone DF9 | 1/4000 |
| HSP70 | WEHI mAb Lab | clone N6 | 1/5000 |
| ACTIN | Sigma-Aldrich | #A2228 clone AC-74 | 1/5000 |
| goat anti-rabbit | Southern Biotech | #4010-05 | 1/5000 |
| goat anti-mouse | Jackson Immuno Research | #115-035-071 | 1/5000 |
| goat anti-rat | Southern Biotech | #3010-05 | 1/5000 |
| goat anti-hamster | Southern Biotech | #6060-05 | 1/1000 |

### Fractionation of Nuclear and Cytoplasmic Proteins

To determine the sub-cellular localization of full-length and truncated MNT proteins after CRISPR/Cas9, 3x10^6^ cells of WT and *MNT KO* BL-2 and *Eμ-Myc* lymphoma cells were lysed in 100 μL of Cytoplasmic Lysis Buffer (CLB) (10 mM Hepes, pH 7.5, 10 mM KCl, 2 mM MgCl_2_, 0.1 mM EDTA, 0.2% IGEPAL CA-630 (Merck, Cat#9002-93-1), 1 mM PMSF and protease inhibitor cocktail (Roche, Cat#5892970001) on ice for 10 min. After centrifugation at 1200 rpm for 5 min, the supernatant (cytoplasmic fraction) was collected into a clean 1.5 mL tube. The nuclear pellet was washed three times by gentle resuspension in 400 μL CLB followed by centrifugation at 1200 rpm for 5 min, removal of the supernatant, and resuspension in 100 μL of Nuclear Lysis Buffer (50 mM Tris pH 8.0, 150 mM NaCl, 0.1% SDS, 1% IGEPAL CA-630, 1 mM PMSF and protease inhibitor cocktail) and incubated on ice for 15 min. After centrifugation at 13,000 rpm for 15 min, the supernatant (nuclear fraction) was collected into a clean 1.5 mL tube. The cytoplasmic and nuclear fractions were centrifuged again at 13,000 rpm for 15 min, then 25 μL was analyzed by western blotting.

### IP/Western assay for BIM Complexed to Pro-Survival BCL-2 Family Proteins

Parental and BCL-2^hi^ *Mnt^fl/fl^ Eμ-Myc/CreERT2* lymphoma cells (7x10^6^) treated for 12 hr with DMSO or ABT-199 (Venetoclax) (1 and 5 μM) were lysed in 1 mL of IP lysis buffer (20 mM Hepes pH 7.5, 200 mM NaCl, 5% Glycerol, 1% IGEPAL CA-630, 1 mM PMSF and proteases cocktail) and incubated on ice for 20 min. Supernatants were collected after centrifugation at 10,000 rpm for 15 min at 4°C and 35 μL (∼3%) was kept for Input control. Supernatants were then incubated with 1 μg BIM antibody or Ig control overnight at 4° C on a rotating wheel. Then, 30 μL of protein G magnetic beads (Invitrogen, Cat#10003D), pre-washed with IP lysis buffer, was added to each sample and incubated for 2 hr on a rotating wheel at 4°C. Samples were then placed on a magnet rack for 2 min and supernatants collected for “unbound” control. Following washing of the beads with 1 mL IP lysis buffer for 5 min (x2) on a rotating wheel, immunoprecipitated proteins were eluted by heating with 60 μL elution buffer (0.1% SDS, 1 mM EDTA, 1X NuPAGE LDS sample buffer (Invitrogen, Cat#NP0007) at 95° C for 5 min. Aliquots (30 μL) of input, unbound and IP sample were analyzed by western blotting.

### Cell Competition Assays

*Mnt^fl/fl^ Eμ-Myc/CreERT2* lymphoma cells infected with *MCL-1/GFP* retrovirus were treated with 4-OHT (0.5 μM) or medium alone (control) as above and harvested on d4. Then, 2x10^5^ viable 4-OHT-treated cells (*Mnt* ko, GFP^+^) were mixed (1:1 or 2:1) with their uninfected parental cells (*Mnt*^+/+^, GFP^-^) and cultured at 37° C in 5 mL fresh DMEM medium lacking 4-OHT. In parallel, control cells (*Mnt^fl/fl^*, GFP^+^) were cultured 1:1 with the same parental CL (*Mnt*^+/+^, GFP^-^). The relative proportions of viable GFP-negative and GFP-positive cells in the mixed cultures were determined on days 0, 2 and 4 by FACS (Fortessa BD Biosciences). The same protocol was used for the control *MCL-1^hi^ Mnt^+/+^ Eμ-Myc/CreERT2* CL.

For *BaxBak dko* clones of *Mnt^fl/fl^ Eμ-Myc/CreERT2* lymphoma CLs, 2x10^5^ cells were incubated at 37° C with 4-OHT (0.5 μM) or medium alone for 20 hr, then diluted 8-fold with fresh DMEM medium and cultured until d4. Then, 2x10^5^ viable 4-OHT-treated cells (*Mnt^-/-^BaxBak dko*, BFP^+^) were mixed (1:1) with the parental CL (*Mnt^fl/fl^ Bax^+/+^Bak^+/+^,* BFP^-^) and cultured at 37° C in 5 mL fresh DMEM lacking 4-OHT. In parallel, control cells (*Mnt^fl/fl^ BaxBak dko,* BFP^+^) were mixed 1:1 with the parental CL (*Mnt^fl/fl^ Bax^+/+^Bak^+/+^,* BFP^-^) and cultured. The relative proportions of viable BFP-negative and BFP-positive cells in the mixed cultures were determined by FACS at 0, 48 and 96 hr.

For *MNT KO* clones of BL lines, 2x10^5^ viable (mCherry^+^ GFP^+^) cells were mixed (1:1) with the Cas9^+^ *MNT^+/+^* parental CL (mCherry^+^ GFP^-^) and cultured in 5 mL fresh medium. For the control, two independent doxycycline-untreated *MNT^+/+^*sgRNA (mCherry^+^ GFP^+^) cells were cultured 1:1 with the same parental CL and the relative proportions of viable GFP-negative and GFP-positive cells were determined by FACS on d0 and d4.

### Cell Cycle Analysis

*Mnt^fl/fl^ Eμ-Myc/CreERT2* lymphoma cells overexpressing hMCL-1 were treated with 4-OHT or medium alone as described above, centrifuged on d4, washed with medium lacking 4-OHT and then 2x10^6^ viable cells were incubated in 4 mL medium containing 10 μM BrdU (Invitrogen, Cat #00-4440-51A) for 3 hr at 37°C. The cell cycle profile was determined using a BrdU staining Kit for Flow Cytometry PE (Invitrogen, Cat #8812-6600) as described by the manufacturer.

### Intracellular Antibody-Staining for Flow Cytometry

After treatment with 4-OHT or medium alone (control), cells were collected on d4 and washed with PBS, then fixed and permeabilized in 1 mL PFA buffer (0.5% paraformaldehyde, 0.2% Tween-20, 0.1% BSA in PBS) overnight at 4°C. Cells were pelleted, washed with 2 mL PBA buffer (PBS containing 0.1% BSA), resuspended in 50 μL PBA containing antibody (see below) and incubated on ice for 60 min. Cells were then washed with 2 mL PBA buffer and resuspended in 50 μL PBA containing goat anti-rabbit IgG antibody conjugated to Alexa Fluor 647 (dilution 1:1000, Life Technologies, Cat #A21244) for 30 min on ice. After pelleting at 1500 rpm and washing with 2 mL PBA, cells were analyzed using a Fortessa Flow cytometer (BD Biosciences). Antibodies used were against c-MYC (dilution 1:1500, Cell Signalling, clone D84C12, Cat #9718), H3K9m3 (dilution 1:3000, Abcam, Cat #ab8898) and monoclonal rabbit IgG isotype control (dilution 1:1500, Cell Signalling, clone DA1E, Cat #05/2018). For each condition, the median fluorescence intensity (MFI) values of c-MYC and H3K9me3 were determined by FlowJo. Data are presented as the fold change of the levels of c-MYC and H3K9me3 determined by normalizing MFI of *Mnt ko* cells to control cells.

### Cell Senescence Flow Cytometry Assay (SA-β-Gal Staining)

*BaxBak dko Mnt^fl/fl^*/*Eμ-Myc/CreERT2* cells (2x10^5)^ were incubated with 4-OHT (0.5 μM) or medium alone for 20 hr, diluted 8-fold with fresh medium and cultured at 37° C until d4. Then, 5x10^5^ viable cells were cultured in 4 mL fresh medium for a further 24 hr. Senescent ( β -galactosidase-positive) cells were detected by using the Senescence Green Flow Cytometry Assay kit (Thermo Fisher, Cat #C10840) as described by the manufacturer.

### Drug Sensitivity Assays

*hMCL-1/GFP*, *hBCL-2/GFP* or *GFP* virus-infected cells harvested on d4 after treatment with 4-OHT or medium alone, were plated in duplicate at 3x10^4^ cells/well in flat-bottom 96-well plates in 60 μL DMEM. Drugs were then added at the indicated final concentrations and the cells were incubated at 37° C for 24 hr.

Cas9^+^ parental and *MNT KO* Burkitt Lymphoma cells were seeded in duplicate at 3x10^4^ cells/well in a flat-bottom 96-well plate in 60 μL RPMI. Drugs were then added at the indicated final concentrations and the cells were incubated at 37° C for 24 hr and 48 hr.

Viable cells (PI^-^ or DAPI^-^) were quantified by flow cytometry (Fortessa - BD Biosciences) and data analysed by Flowjo v10.9. Viability was calculated relative to DMSO-treated control. All data are presented as mean ± SEM of at least 3 independent experiments.

Drugs used were: S63845 (MCL-1 inhibitor, MedChemExpress, Cat #HY100741), ABT-199 (Venetoclax, MedChemExpress, Cat #HY-15531), Doxorubicin (MedChemExpress, Cat #HY15142), Cisplatin (MedChemExpress, Cat #HY17394), Vincristine (Sigma-Aldrich, Cat #V8879), Nutlin-3a (Cayman Chemicals, Cat #18585), Ionomycin (Sigma-Aldrich, Cat #I9657) and Etoposide (MedChemExpress, Cat #HY13629).

### Analysis of Cut&Run Data Obtained by Mathsyaraja et al (37)

CUT&RUN data from *wt* and *Max^-/-^* B220^+^ spleen cells were obtained from 9 week old mice (GSM3897577-GSM3897578, GSM3897581-GSM3897584). Analysis was performed as previously described (Feng Yan et al, Genome Biology 2020). Briefly, FASTQ files were quality-checked with FastQC (0.11.8) and trimmomatic (0.39) used to remove low-quality reads and bases, and adapter sequences. Trimmed FASTQ files were then aligned to the mouse genome (mm10) with bwa-mem (0.7.17-r1188), and low-quality (<Q30), duplicated and unmapped reads were removed. BAM files were converted to bigwig files using deepTools (3.5.0) with reads per genomic content normalization (scaled to achieve average 1x genomic coverage) and then uploaded to the UCSC genome browser for visualization.

### Statistical Analysis

Statistical comparisons were made using unpaired two-tailed Student’s *t-test* with Prism v10 software (GraphPad, San Diego, CA, USA). Data are shown as means ± SEM (Standard Error of the Mean) with P ≤ 0.05 considered statistically significant. Mouse survival analysis was carried out using GraphPad Prism (Version 10) and significance determined using log-rank (Mantel–Cox) test.

## Data Availability Statement

Renewable materials and protocols are available without unreasonable restrictions from the first or corresponding author.

## Authors’ Contributions

**H.V.Nguyen**: Conceptualization, design and performance of most experiments, co-supervision of M.M., data assembly, writing-review and editing. **M. Michla**: performed CRISPR/Cas9 *MNT* deletion for Burkitt Lymphoma cells. **F. Yan**: analysis of published CUT&RUN data. **N.M.Davidson**: Review of bioinformatics analysis. **G.L.Kelly**: Advice on design and interpretation of experimental data, writing -review, resources. **A.Strasser**: Advice on design and interpretation of experimental data, writing -review, resources. **S.Cory**: Conceptualization, resources, supervision, data curation, funding acquisition, data assembly, writing-original draft, revised drafts, project administration. All authors reviewed and approved the final manuscript.

## Acknowledgements

The authors thank Giovanni Siciliano and his team in WEHI BioServices for help with the animal experiments and husbandry; Simon Monard and his team at WEHI’s cell sorting facility; and Peter Maltezos with skilled assistance in preparation of the figures.

This work was supported by grants and fellowships from the Australian National Health and Medical Research Council (Program Grant 1016701 (S.C.); Investigator Grant 1020363 (A.S.); Investigator Grant 2034395 (G.L.K.); Investigator Grant GNT2016547 (N.M.D.); Cancer Council Victoria Grant-in-Aid to S.C. and G.L.K.; CASS Foundation grant (9289) to H.V.N.; philanthropic support to WEHI, and infrastructure grants to WEHI through the Victorian State Government Operational Infrastructure Support and the Australian Government NHMRC Independent Research Institute Infrastructure Support Schemes.

