## Supplemental Table and Figs for "MNT loss in MYC-driven B lymphoma cells enhances apoptosis, inhibits proliferation and increases sensitivity to cancer drugs"

**Table S1 Characterisation of *Mnt* deletable and control *Eμ-Myc/CreERT2* Lymphoma Cell Lines**

| *Mnt* Genotype[1] | 1° Lymphoma[1]<br>mouse #<br>sex<br>age | Cell Line[1]<br>tissue origin<br>immunophenotype<br>differentiation stage | Nutlin-3a sensitivity[2] | *p53* status[2] | Transplantation[3]<br>mouse #s, sex | Treatment[3] | Survival<br>(d) |
|---|---|---|---|---|---|---|---|
| $Mnt^{+/+}$ | #1549<br>F<br>116 d | spleen<br>$CD19^+CD43^+ IgM^+$<br>B | S | wt | #9737-39 F<br><br>#9740-42 F | vehicle<br><br>tamoxifen | 18,18,18<br><br>20,21,19 |
| | #1194<br>F<br>170 d | thymic mass<br>$CD19^+CD43^+IgM^+$<br>B | S | wt | #9643-45 F<br><br>#9646-48 F | vehicle<br><br>tamoxifen | 20,20,20<br><br>21,21,21 |
| | #1190<br>F<br>301d | spleen<br>$CD19^+CD43^+IgM^-$<br>pro-B | R | mut | #9819-21 F<br><br>#9822-24 F | vehicle<br><br>tamoxifen | 14,14,14<br><br>17,19,16 |
| | #1591<br>M<br>46 d | spleen<br>$CD19^+ CD43^-IgM^-$<br>pre-B | R | mut<br>(normal<br>size pp) | #10346-48 M<br><br>#10349-51M | vehicle<br><br>tamoxifen | 17,19,18<br><br>20,19,20 |
| $Mnt^{fl/fl}$ | #1271<br>F<br>90 d | thymic mass<br>$CD19^+CD43^-IgM^+$<br>B | S | wt | #9743-45 F<br><br>#9746-48 F | vehicle<br><br>tamoxifen | 28,32,18*<br><br>42,42,154* |
| | #1721<br>F<br>230 d | LNs<br>$CD19^+ CD43^+ IgM^+$<br>B | S | wt | #9853-55 F<br><br>#9856-58 F | vehicle<br><br>tamoxifen | 21,20,21<br><br>36,40,36 |
| | #1506<br>F<br>85 d | spleen<br>$CD19^+ CD43^+ IgM^+$<br>B | S | wt | #4859-61 F<br><br>#9862-64 F | vehicle<br><br>tamoxifen | 13,13,13<br><br>19,19,17 |

| | | | | | | | |
|---|---|---|---|---|---|---|---|
| | #2221<br>M<br>72 d | spleen<br>$CD19^{+}$ $CD43^{+}$ $IgM^{-}$<br>pro-B | S | wt | #10292-94 M<br><br>#10295-97 M | vehicle<br><br>tamoxifen | 32,32,53<br><br>127,167,167* |
| | #654<br>F<br>79 d | thymic mass<br>$CD19^{+}CD43^{-}IgM^{-}$<br>pre-B | S | wt | #9776-78 F<br><br>#9779-81 F | vehicle<br><br>tamoxifen | 17,17,17<br><br>30,34,27 |
| | #760<br>F<br>68 d | spleen<br>$CD19^{+}CD43^{-}IgM^{-}$<br>pre-B | S | wt | #9703-9705<br><br>#9706-9708 | vehicle<br><br>tamoxifen | 14,9,14<br><br>26,27,27 |
| | #799<br>F<br>67 d | thymic mass<br>$CD19^{+}CD43^{-}IgM^{-}$<br>pre-B | S | wt | #9788-90 F<br><br>#9791-93 F | vehicle<br><br>tamoxifen | 21,21,21<br><br>35,35,35 |
| | #1129<br>F<br>155 d | Spleen<br>$CD19^{+}CD43^{-}IgM^{-}$<br>pre-B | S | wt | #9631-33 F<br><br>#9634-36 F | vehicle<br><br>tamoxifen | 14,14,14<br><br>45,335*,335* |
| *Mnt*$^{fl/fl}$ | #643<br>F<br>364 d | mesLN<br>$CD19^{+}CD43^{-}IgM^{+}$<br>B | R | mut<br>(small<br>pp) | #9731-33 F<br><br>#9734-36 F | vehicle<br><br>tamoxifen | 18,26,25<br><br>35,35,35 |
| | #758<br>F<br>128 d | spleen<br>$CD19^{+}CD43^{-}IgM^{+/-}$<br>pre-B/B | R | mut | #9725-27 F<br><br>#9728-30 F | vehicle<br><br>tamoxifen | 18,18,18<br><br>25,25,25 |
| | #1344<br>F<br>219 d | spleen<br>$CD19^{+}CD43^{+}IgM^{+}$<br>B | R | mut<br>(no<br>pp) | #9785-87 F<br>#9782-84 F<br><br>#9791-93 F<br>#9785-87 F | vehicle<br><br><br>tamoxifen | 14,14,14<br>13,13,13<br><br>19,21,20<br>18,18,18 |

| | | | | | | | |
|---|---|---|---|---|---|---|---|
| | #2297<br>F<br>323 d | spleen<br>$CD19^+CD43^+IgM^+$<br>B | R | mut | #10334-36 F<br><br>#10337-39 F | vehicle<br><br>tamoxifen | 22*,24, 26<br><br>24,53,40* |
| | #2301<br>F<br>89 d | spleen<br>$CD19^+CD43^+IgM^+$<br>B | R | mut | #10299-300 F<br><br>#10301-03 F | | 15,15,15<br><br>23,22,23 |
| | #691<br>F<br>112 d | thymus<br>$CD19^+CD43^{+/-}IgM^-$<br>pre-B | R | mut | ns | ns | ns |

[1]*Lymphoma cell lines (CLs) were established from indicated lymphomatous tissue from autopsied* $Mnt^{+/+}$ *and* $Mnt^{fl/fl}$ *Eμ-Myc/CreERT2 mice, as described in text and Methods. Cell surface immunophenotypes were determined by flow cytometry after staining with the following antibodies: rat anti-mouse CD19-PerCP/Cyanine5.5 (BioLegend, clone 1D3/CD19, Cat # 152405, dilution 1:300) , rat anti-mouse CD43-PE (BD Biosciences, clone S7, Cat #553271, dilution 1:300), monoclonal anti-mouse CD54.1-APC (BioLegend, clone A20, Cat #110713, dilution 1:300), anti-mouse IgM-FITC (WEHI mAb lab, clone 5-1, dilution 1:400)*

[2, 3] *p53 status was determined as described in Methods; p53 wt cells die when treated with Nutlin 3a, but p53 mutant cells remain viable.*

[4] *Multiple mice transplanted with individual lymphoma CLs were treated with either vehicle or tamoxifen as described in text and their health was regularly monitored for >100 d, until the experiment was terminated on the day indicated. Mice deemed sick by independent, trained technicians were euthanized and autopsied for signs of lymphoma (enlarged organs, abnormal blood profile).*

** indicates transplant recipients that were censored from survival curves because they had no detectable leukemia or lymphoma when autopsied.*

*ns indicates transplantation experiment failed due to technical reasons*

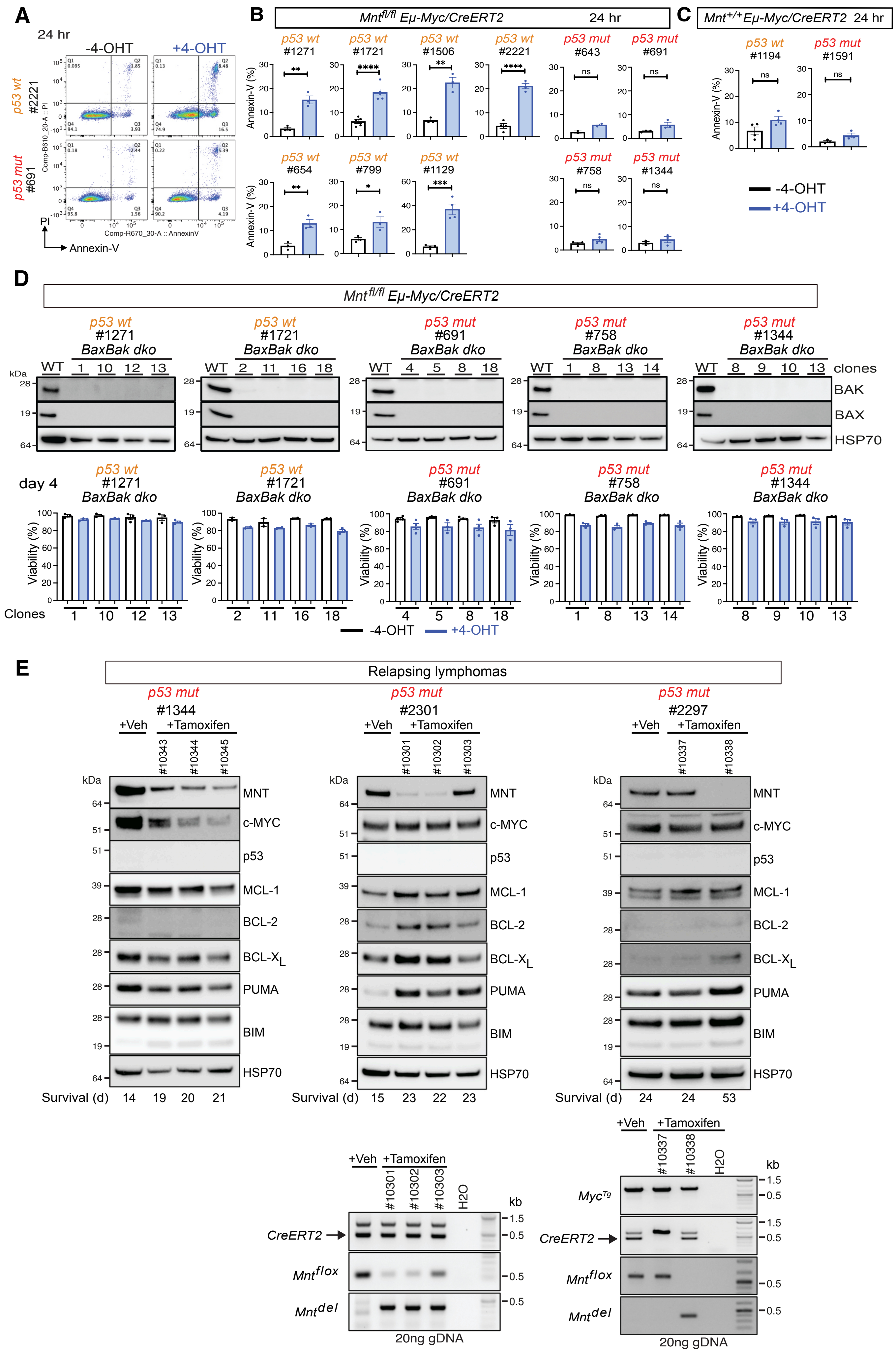
A
24 hr
-4-OHT
+4-OHT
p53 wt #2221
p53 mut #691
PI
Annexin-V
B
Mnt fl/fl Eμ-Myc/CreERT2 24 hr
p53 wt #1271
p53 wt #1721
p53 wt #1506
p53 wt #2221
p53 mut #643
p53 mut #691
p53 wt #654
p53 wt #799
p53 wt #1129
p53 mut #758
p53 mut #1344
Annexin-V (%)
C
Mnt +/+ Eμ-Myc/CreERT2 24 hr
p53 wt #1194
p53 mut #1591
-4-OHT
+4-OHT
D
Mnt fl/fl Eμ-Myc/CreERT2
p53 wt #1271 BaxBak dko
p53 wt #1721 BaxBak dko
p53 mut #691 BaxBak dko
p53 mut #758 BaxBak dko
p53 mut #1344 BaxBak dko
clones
BAK
BAX
HSP70
day 4
Viability (%)
Clones
-4-OHT
+4-OHT
E
Relapsing lymphomas
p53 mut #1344
p53 mut #2301
p53 mut #2297
+Veh
+Tamoxifen
MNT
c-MYC
p53
MCL-1
BCL-2
BCL-XL
PUMA
BIM
HSP70
Survival (d) 14 19 20 21
Survival (d) 15 23 22 23
Survival (d) 24 24 53
Myc Tg
CreERT2
Mnt flox
Mnt del
20ng gDNA

**Figure S1.** *Mnt* deletion in *Mnt$^{fl/fl}$ Eμ-Myc/CreERT2* lymphoma cells**. A-C**, Apoptosis triggered by *Mnt* deletion starts earlier in *p53 wt* than *p53 mut* lymphoma CLs. **A**, Typical flow cytometric analysis of cell viability, performed by staining with Annexin-V and PI at 24 hr . **B-C,** Annexin-V staining of *p53 wt* and *p53 mut Mnt$^{fl/fl}$* (**B**) and control *Mnt$^{+/+}$*(**C**) *Eμ-Myc/CreERT2* lymphoma CLs. Lymphoma cells were incubated for 20 hr in medium containing 0.5 μM 4-OHT (blue columns) or medium alone (open columns) then diluted 8-fold with fresh medium. Cell aliquots taken at 24 hr were stained with Annexin-V and PI, then analysed by flow cytometry (**A**). Bar graphs show % Annexin-V$^+$ (non-viable) cells in at least 3 independent experiments (dots), shown as mean $\pm$ SEM; *P$\leq$ 0.05, **P$\leq$ 0.01, *** P$\leq$ 0.001, ****P$\leq$ 0.0001. **D**, Cell death induced by *Mnt* loss is BAX/BAK dependent. Upper panels show western blots of *BaxBak dko* clones derived by CRISPR/Cas9 from the indicated parental *p53 wt* and *p53 mut Eμ-Myc/CreERT2* lymphoma CLs (see Methods). Lower panels compare the viability on day 4 of clones treated with 4-OHT (blue columns) or medium alone (open columns), determined by flow cytometry after staining with PI (see Fig S6E and Methods). Results are shown for at least 3 independent experiments (dots) and expressed as mean $\pm$ SEM. **E,** Western blot analysis of relapsing lymphomas from tamoxifen-treated mice that had been transplanted with *p53 mut Mnt$^{fl/fl}$ Eμ-Myc/CreERT2* lymphoma CLs (see text). Western blots are shown above, PCR analyses below. Each lymphoma CL was injected into 6 recipients, identified by #s immediately above blots (see also Table S1).

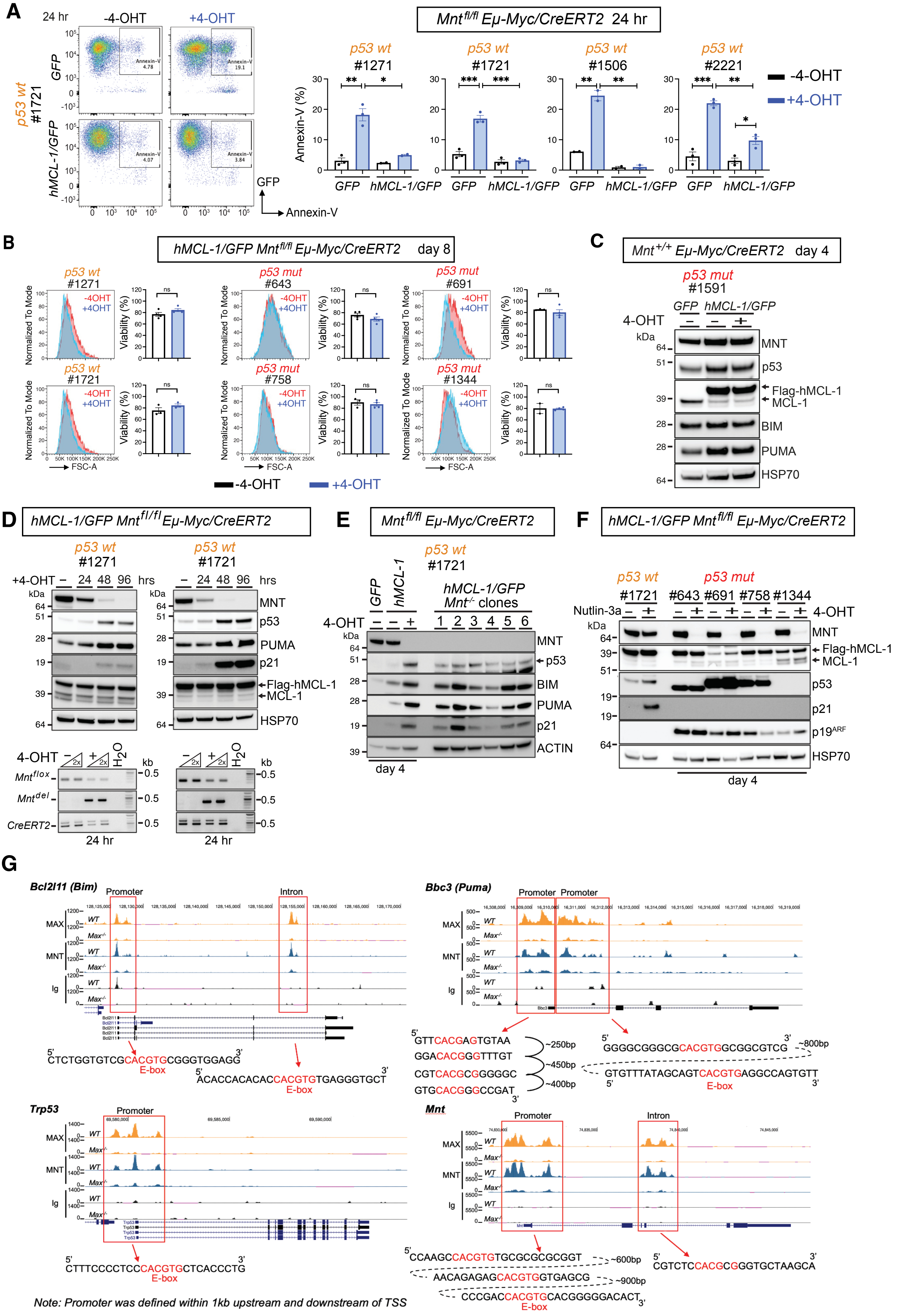

Nguyen et al Figure S2

**Figure S2.** Analysis of cell death signals triggered by *Mnt* deletion in *Eμ-Myc* lymphoma cells. **A**, **B,** Over-expression of MCL-1 protects p53 *wt* and p53 *mut Eμ-Myc/CreERT2* lymphoma cells from apoptosis induced by *Mnt* deletion. *Eμ-Myc/CreERT2* lymphoma CLs expressing *hMCL-1/GFP* or control *GFP* retrovirus were incubated for 20 hr +/- 4-OHT then cultured in fresh medium lacking 4-OHT as shown in Fig 2A. **A,** Left panel shows a typical FACS analysis of cells stained with Annexin-V at 24 hr and right panel summarizes Annexin-V staining for four p5*3 wt* CLs treated with 4-OHT (blue columns) or medium alone (empty columns). Results are shown for 2-3 independent experiments (dots) performed with each indicated CL, expressed as mean $\pm$ SEM; *P$\leq$ 0.05, **P$\leq$ 0.01, *** P$\leq$ 0.001. **B**, Flow cytometry performed on *p53 wt* and *p53 mut* $Mnt^{fl/fl}$ *Eμ-Myc/CreERT2* CLs on day 8 showed little change in cell size (left panels) or viability (right panels), determined by PI staining, between cells that had been treated earlier with 4-OHT (blue columns) or medium alone (open columns). Results are shown for 2-3 independent experiments (dots) and expressed as mean $\pm$ SEM; ns=not significant. **C**, 4-OHT-treatment did not alter the level of p53, BIM or PUMA in control *p53 mut* $Mnt^{+/+}$ Eμ-*Myc/CreERT2* lymphoma CL (#1591) infected with *hMCL-1/GFP* virus. **D**, Kinetics of *Mnt* deletion, *p53* induction, and expression of PUMA and p21 protein. Top panel shows western blot of two *p53 wt* $MCL\text{-}1^{hi}$ $Mnt^{fl/fl}$ *Eμ-Myc* CLs (#1271 and #1721) analysed 0, 24, 48 and 96 hr after 4-OHT treatment for 20 hr. Lower panels show PCR analysis of *Mnt* deletion ($Mnt^{del}$) 24 hr after 4-OHT treatment. **E**, p53, BIM, PUMA and p21 protein levels remain elevated in $Mnt^{-/-}$ clones isolated 8 days after 4-OHT-treatment of *p53 wt* $MCL\text{-}1^{hi}$ $Mnt^{fl/fl}$ *Eμ-Myc/CreERT2* lymphoma CL (#1721) (see protocol in Fig. S3B). Control in first three tracks shows western blot of lysates prepared on day 4 from cells that had been treated on day 1with 4-OHT. **F**, Western blot analysis showing that p21 protein is undetectable in MCL-1 over-expressing *p53 mut* $Mnt^{fl/fl}$ *Eμ-Myc/CreERT2* lymphoma CLs on day 4 after brief 4-OHT treatment. Nutlin-3a (5μM) treatment of *p53 wt* $MCL\text{-}1^{hi}$ *Eμ-Myc* CL (#1721) (first two tracks) provides a positive control for p21 induction. p19ARF protein is detectable in *p53 mut* but not *p53 wt* Eμ-*Myc* lymphoma cells. **G**, MNT and MAX binding sites in *Bcl2l11*(*Bim*), *Bbc3* (*Puma*), *p53 (Trp53)* and *Mnt* gene loci, compared to a non-specific control, Ig. Binding was determined by analysis of CUT&RUN data sets obtained by Mathsyaraja et al {Mathsyaraja, 2019 #19548} from $B220^{+}$ splenic B cells of WT and $Max^{-/-}$ 9 week-old mice having a mixed 129/C57BL/6 background.

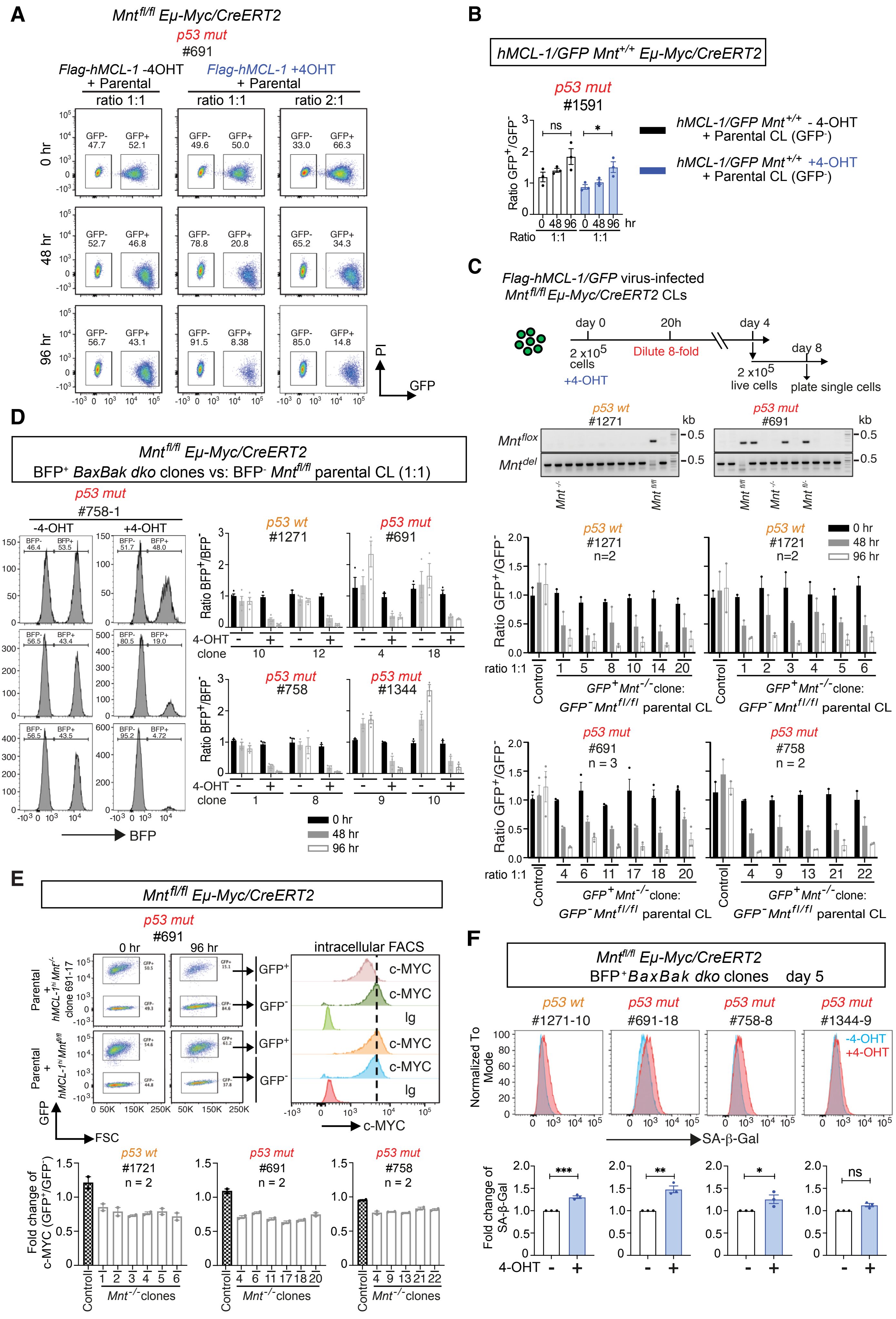

Nguyen et al Figure S3

**Figure S3**. *Mnt* deletion reduces cell proliferation in apoptosis-resistant (*MCL-1$^{hi}$* or *BaxBak dko) Mnt$^{fl/fl}$ Eμ-Myc/CreERT2* lymphoma cells. **A**, Typical flow cytometric analysis of cell competition assay described in Fig 3B. **B,** Cell competition assay showing that 4-OHT treatment does not alter proliferation of control *p53 mut MCL-1$^{hi}$ Mnt$^{+/+}$ Eμ-Myc/CreERT2* cell line (#1591). The bar graph shows GFP$^{+}$/GFP$^{-}$ ratios over time determined by flow cytometry. **C,** *Mnt$^{-/-}$* clones of *p53 wt* and *p53 mut Eμ-Myc/CreERT2* CLs infected with h*MCl-1/GFP* virus have a severe proliferative defect. Top panel shows protocol and central panel shows PCR genotype analysis of clones isolated on day 8 following brief 4-OHT treatment of indicated lymphoma CLs. Examples of *Mnt$^{fl/fl}$*, *Mnt$^{fl/-}$* and *Mnt$^{-/-}$* genotypes are indicated. Bottom panels show results of competition proliferation assays performed between the indicated *Mnt$^{-/-}$* (GFP$^{+}$) clones and *Mnt$^{fl/fl}$* (GFP$^{-}$) parental cells mixed (1:1). The relative proportion of GFP$^{+}$ and GFP$^{-}$ cells was determined at 0, 48 and 96 hr by flow cytometry. The diminishing GFP$^{+}$/GFP$^{-}$ ratio over time indicates that the *Mnt$^{-/-}$* clones had a proliferative defect. n= no of independent experiments performed. **D,** *Mnt*-deleted clones of *p53 wt* and *p53 mut BaxBak dko Eμ-Myc/CreERT2* lymphoma CLs have a marked proliferative defect. To delete *Mnt*, 2x10$^{5}$ cells from *BaxBak dko* clones generated from *Eμ-Myc/CreERT2* CLs (see Fig S1D) were incubated with 4-OHT (0.5 μM) for 20 hr, then diluted 8-fold with fresh medium and cultured at 37$^{o}$ C until day 4. Then, 2x10$^{5}$ viable cells (BFP$^{+}$) were mixed (1:1) with *Mnt$^{fl/fl}$ Bax$^{+/+}$Bak$^{+/+}$*parental cells (BFP$^{-}$) and cultured at 37$^{o}$C in 5 mL fresh medium. In parallel, control untreated *BaxBak dko Mnt$^{fl/fl}$ Eμ-Myc* clones (BFP$^{+}$) were cultured in competition, mixed 1:1 with the *Bax$^{+/+}$Bak$^{+/+}$* parental CL (BFP$^{-}$). The relative proportion of viable BFP$^{+}$/BFP$^{-}$ cells in the mixed cultures was determined by flow cytometry at 0, 48 and 96 hr. The left panels show typical flow cytometric analysis and the right panels show the diminishing BFP$^{+}$/BFP$^{-}$ ratios over time for two clones of each of the 4 indicated lymphoma cell lines. **E,** Decreased level of MYC in *Mnt*-deleted clones isolated from *p53* wt and *p53 mut MCL-1$^{hi}$ Eμ-Myc/CreERT2* lymphoma CLs. Cells taken from competition assay (Fig. S3C) at 96 hr were permeabilized, stained with c-MYC or Ig antibody followed by intracellular flow cytometry. Typical flow cytometric analysis (top panels) shows MYC level at 96 hr. The bottom panels show fold decrease in median fluorescence of MYC in GFP$^{+}$ *MCL-1$^{hi}$ Mnt$^{-/-}$ Eμ-Myc* clones compared to their GFP$^{-}$ parental cells (open columns). In contrast, MYC levels were comparable in the GFP$^{-}$ and GFP$^{+}$ cell populations of parental vs untreated *MCL-1$^{hi}$ Mnt$^{fl/fl}$* cells (hatched columns). n=no of independent experiments performed/clone. **F**, *Mnt* deletion in apoptosis-resistant *BaxBak dko Eμ-Myc* lymphoma cells results in senescence. Clones of indicated *p53 wt* and *p53 mut BaxBak dko Mnt$^{fl/fl}$ Eμ-Myc/CreERT2* lymphoma CLs were treated with 4-OHT (pink) or medium alone (blue) followed by dilution at 20 hr, then permeabilized, stained for SA-β-Gal and analyzed on day 5 by flow cytometry (see Methods). Bottom panels show fold change in median fluorescence of SA-β-Gal in *Eμ-Myc/CreERT2* cells treated with 4-OHT (blue columns) versus medium alone (open columns). Results are shown for at least 3 independent experiments (dots) and expressed as mean ± SEM; *P≤ 0.05, **P≤ 0.01, and ns= not significant.

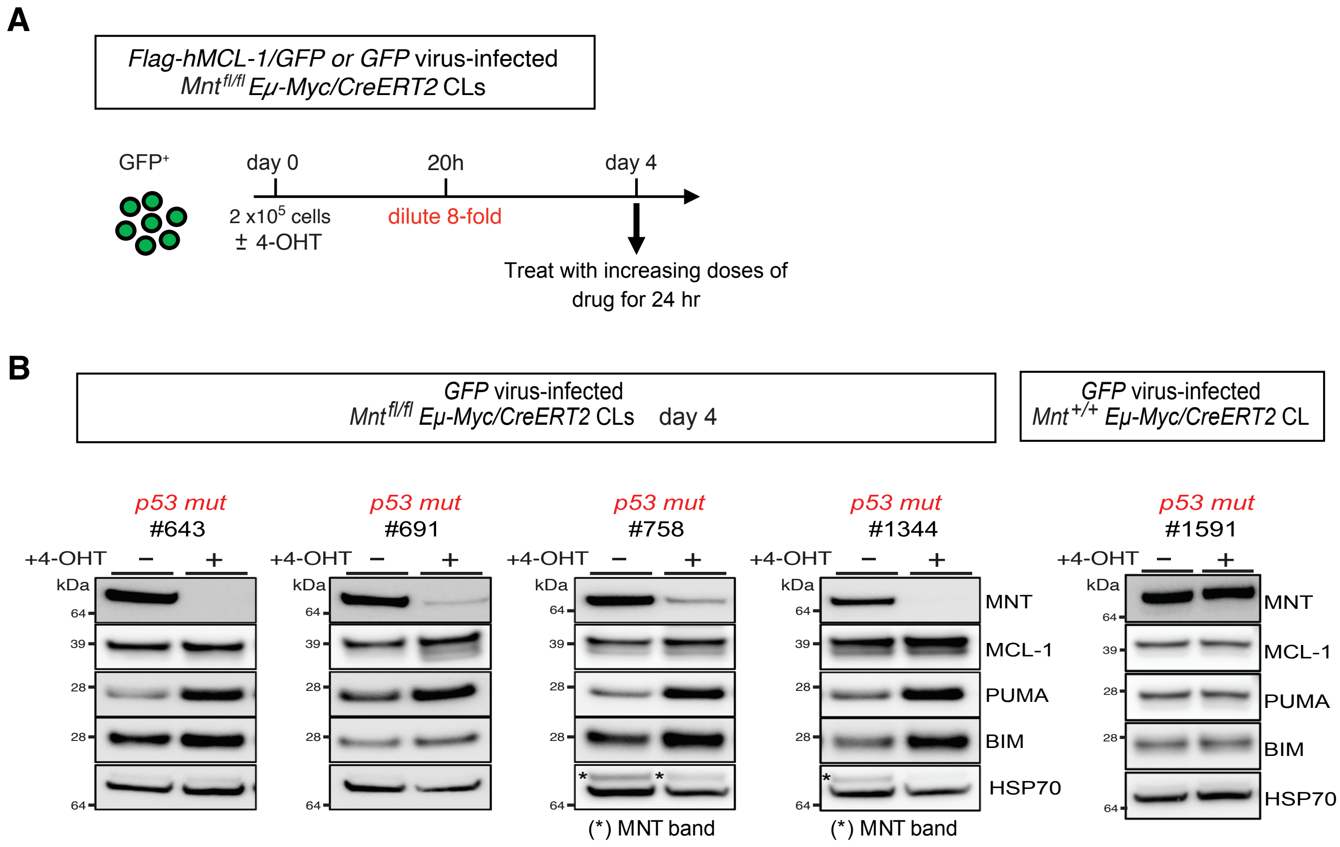
A
Flag-hMCL-1/GFP or GFP virus-infected
Mnt fl/fl Eμ-Myc/CreERT2 CLs
GFP+
day 0
20h
day 4
2 x10^5 cells
± 4-OHT
dilute 8-fold
Treat with increasing doses of drug for 24 hr
B
GFP virus-infected
Mnt fl/fl Eμ-Myc/CreERT2 CLs day 4
GFP virus-infected
Mnt +/+ Eμ-Myc/CreERT2 CL
p53 mut
#643
p53 mut
#691
p53 mut
#758
p53 mut
#1344
p53 mut
#1591
+4-OHT − +
kDa
64
39
28
MNT
MCL-1
PUMA
BIM
HSP70
(*) MNT band

**Figure S4**. MNT loss sensitizes *Eμ-Myc* lymphoma cells to cancer chemotherapeutic drugs. **A,** Protocol for treating *Eμ-Myc* lymphoma CLs with cancer drugs following brief treatment with 4-OHT to delete *Mnt*. 2x$10^5$ *hMCL-1/GFP* or *GFP*-virus infected *Mnt*$^{fl/fl}$ *Eμ-Myc/CreERT2* lymphoma cells were treated with 4-OHT (0.5 μM) or medium alone for 20 hr, then diluted 8-fold. The cells were cultured for a further 3 days and then treated with a range of concentrations of the indicated drug for 24 hr before determining viability by flow cytometry. **B,** Western blot performed on day 4 shows elevated BIM and PUMA protein levels after 4-OHT-mediated *Mnt* deletion in four p53 *mut GFP* virus-infected *Mnt*$^{fl/fl}$ *Eμ-Myc/CreERT2* lymphoma CLs but not in the *Mnt*$^{+/+}$ control CL (#1591). Western blots shown are typical of at least 2 independent experiments for each CL. Comparable western blot data for *Mnt*$^{fl/fl}$ *Eμ-Myc/CreERT2* lymphoma CLs infected with *hMCL-1/GFP* virus-infected is shown in Fig. 2B, C**.**

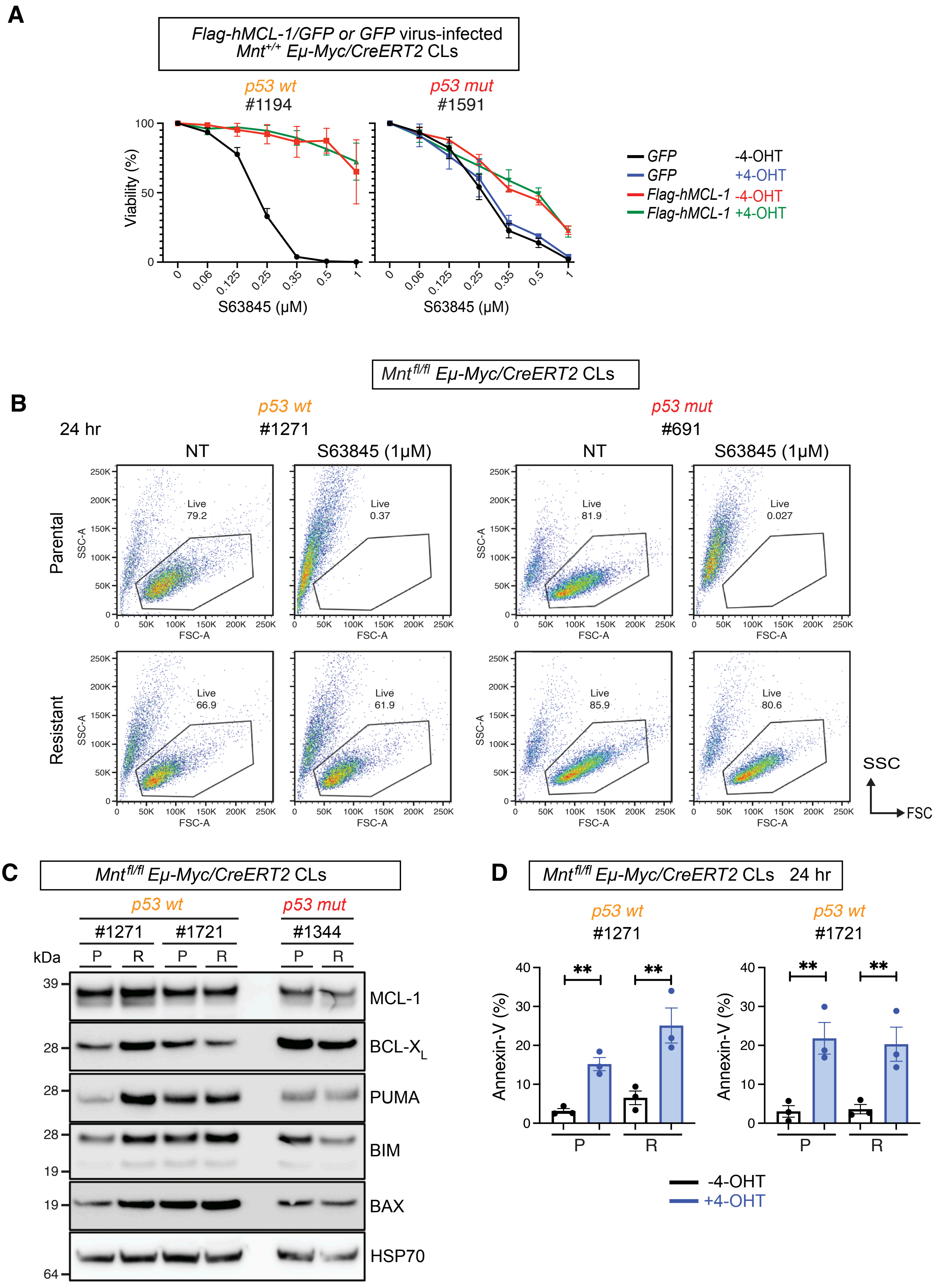

Nguyen et al Figure S5

**Figure S5** Investigating the impact of MNT loss in S63845-resistant *Eμ-Myc/CreERT2* lymphoma cells**. A,** Negative control showing that 4-OHT treatment of indicated *p53 w*t and *p53 mut* $Mnt^{+/+}$ *Eμ-Myc/CreERT2* lymphoma CLs infected with *hMCL-/GFP* or control *GFP* retrovirus has no impact on sensitivity to S63845. **B**, Typical flow cytometric analysis of viability of parental and S63845-resistant $Mnt^{fl/fl}$ *Eμ-Myc/CreERT2* lymphoma cells 24 hr after treatment with S63845 (1 μM). **C,** Expression analysis of the indicated BCL-2 family members in parental (P) and S63845-resistant (R) derivatives of the indicated *p53 wt* and *p53 mut* $Mnt^{fl/fl}$ *Eμ-Myc/CreERT2* lymphoma CLs, determined by western blot analysis. HSP70 expression was used as a loading control. **D,** MNT loss induced by 4-OHT increases apoptosis of both the parental (P) and S63845-resistant (R) derivative of the indicated *p53 wt* $Mnt^{fl/fl}$ *Eμ-Myc/CreERT2* lymphoma CLs. Viability at 24 hr was determined by flow cytometry after staining for Annexin-V and PI. Analysis is shown for 3 independent experiments and plotted as mean ± SEM; *P≤ 0.05, **P≤ 0.01.

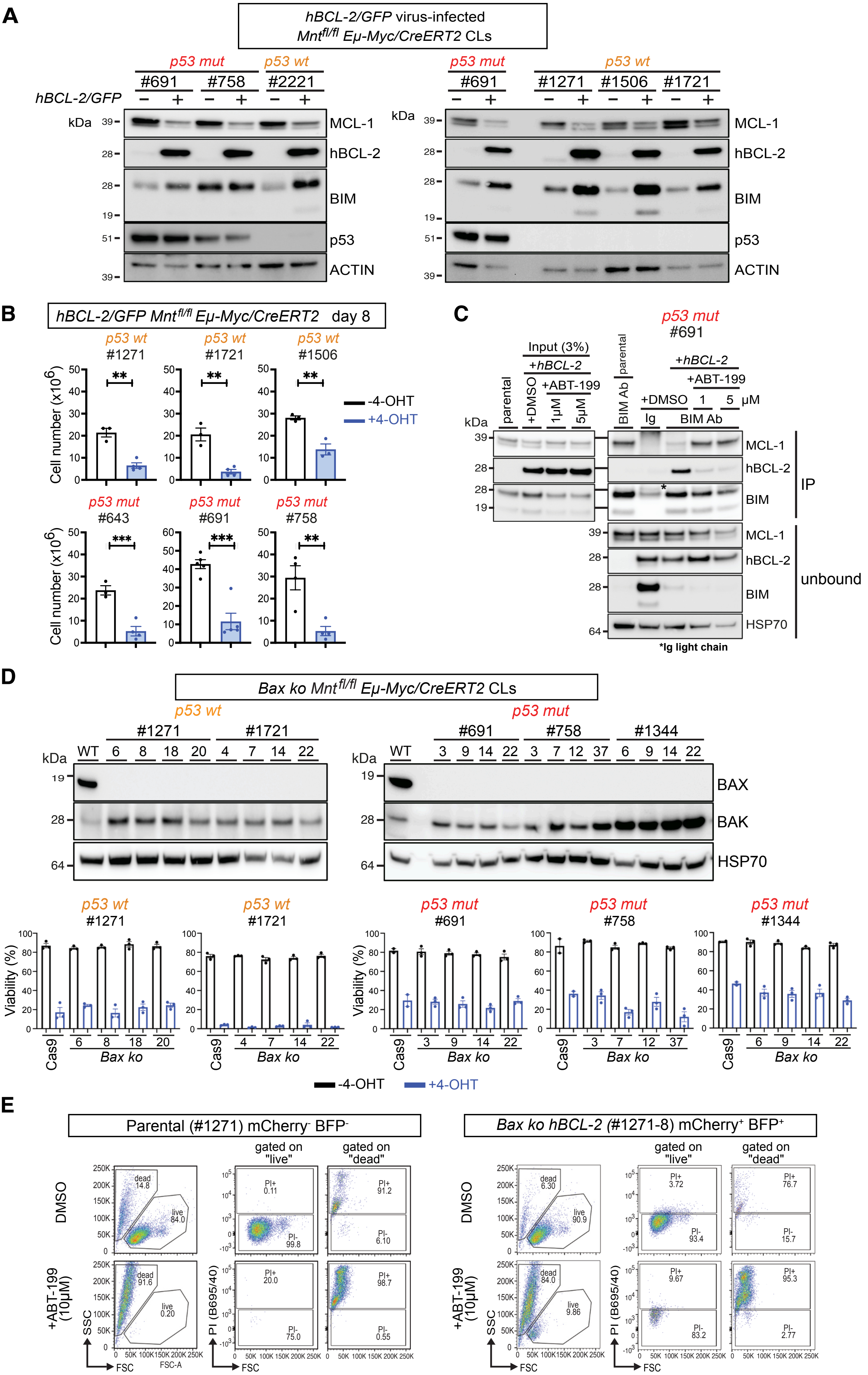

Nguyen et al Figure S6

**Figure S6.** Characterization of *hBCL-2/GFP* virus-infected *Mnt*$^{fl/fl}$ *Eμ-Myc/CreERT2* lymphoma cell lines and their *Bax* knockout (*ko*) derivatives. **A**, Western blot analysis showing that BIM levels increase and MCL-1 levels decrease in both *p53 wt* and *p53 mut Eμ-Myc/CreERT2* lymphoma cells over-expressing BCL-2 compared to parental lymphoma cells. **B,** Reduced proliferation of *p53* wt and *p5*3 *mut* apoptosis-resistant (*BCL-2*$^{hi}$) *Mnt*$^{fl/fl}$ *Eμ-Myc/CreERT2* lymphoma cells following 4-OHT-induced *Mnt* deletion. The indicated h*BCL-2* virus-infected lymphoma CLs were treated +/- 4-OHT, then diluted and cultured as in Fig 2A. Cell number on day 8 was compared for cells treated with 4-OHT (blue columns) versus those treated with medium alone (open columns). Results are shown for at least 3 independent experiments and plotted as mean $\pm$ SEM; *P$\leq$ 0.05, **P$\leq$ 0.01, *** P$\leq$ 0.001. **C**, IP/western blot analysis of *BCL-2*$^{hi}$ *p53 mut Eμ-Myc* lymphoma CL (#691) showing that BCL-2-bound BIM decreased following ABT-199 treatment (1 and 5 μM) for 12 hr, and MCL-1-bound BIM increased (see also Fig 6B). **D,** Characterization of *Bax ko* clones derived from *p53 wt* and *p53 mut Mnt*$^{fl/fl}$ *Eμ-Myc/CreERT2* CLs (see Methods). Western blots in upper panels show the absence of BAX and presence of BAK protein in *Bax ko* clones. HSP70 was used as a loading control. The bar graphs in lower panels show that *Mnt* deletion induced by 4-OHT in *Bax ko* clones results in apoptosis. Independent *Bax ko* clones derived from the indicated *Mnt*$^{fl/fl}$ *Eμ-Myc/CreERT2* lymphoma CLs were treated +/- 4-OHT (0.5 μM) as described in Fig 1A. Cells were collected on day 4, stained with PI and cell viability was determined by flow cytometry. Viability was greatly diminished for *Bax ko* cells that had been incubated with 4-OHT (blue columns) compared to medium alone (open columns), presumably via apoptosis activated via BAK. For the control, parental CLs infected with Cas9 lentivirus was used to assess viability after treatment with 4-OHT. **E,** Typical flow cytometric analysis of ABT-199 treated parental and *Bax ko Eμ-Myc/CreERT2* lymphoma cells (see Fig 6E). The left panels show live and dead cells of parental *Eμ-Myc* lymphoma cells (#1271) (mCherry$^-$ BFP$^-$) and the right panels show analysis of derivative BCL-2$^{hi}$ *Bax ko* clone 8 (mCherry$^+$ BFP$^+$) cells following 24 hr treatment with ABT-199 (10 μM) or DMSO.

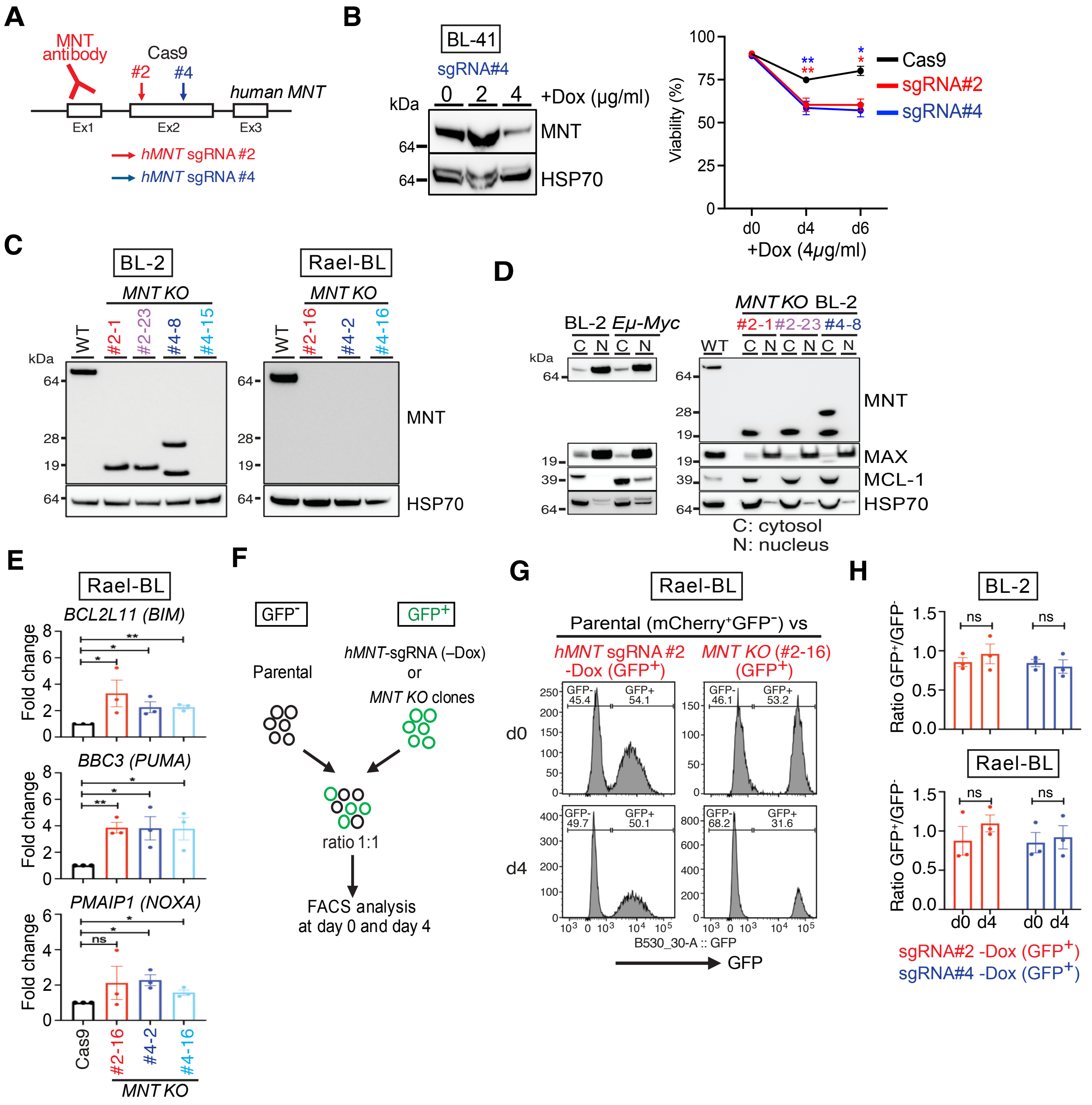
A
MNT antibody
Cas9
#2
#4
human MNT
Ex1
Ex2
Ex3
hMNT sgRNA #2
hMNT sgRNA #4
B
BL-41
sgRNA#4
kDa
0 2 4 +Dox (µg/ml)
MNT
HSP70
Viability (%)
Cas9
sgRNA#2
sgRNA#4
d0 d4 d6
+Dox (4µg/ml)
C
BL-2
Rael-BL
MNT KO
WT #2-1 #2-23 #4-8 #4-15
WT #2-16 #4-2 #4-16
MNT
HSP70
D
BL-2 Eµ-Myc
MNT KO BL-2
#2-1 #2-23 #4-8
C N C N
MNT
MAX
MCL-1
HSP70
C: cytosol
N: nucleus
E
Rael-BL
BCL2L11 (BIM)
BBC3 (PUMA)
PMAIP1 (NOXA)
Fold change
Cas9 #2-16 #4-2 #4-16
MNT KO
F
GFP-
GFP+
Parental
hMNT-sgRNA (–Dox)
or
MNT KO clones
ratio 1:1
FACS analysis
at day 0 and day 4
G
Rael-BL
Parental (mCherry+GFP−) vs
hMNT sgRNA #2 -Dox (GFP+)
MNT KO (#2-16) (GFP+)
d0
d4
GFP- 45.4 GFP+ 54.1
GFP- 46.1 GFP+ 53.2
GFP- 49.7 GFP+ 50.1
GFP- 68.2 GFP+ 31.6
B530_30-A :: GFP
GFP
H
BL-2
Rael-BL
Ratio GFP+/GFP-
ns
d0 d4 d0 d4
sgRNA#2 -Dox (GFP+)
sgRNA#4 -Dox (GFP+)

**Figure S7.** MNT loss in Burkitt Lymphoma cell lines. **A**, Diagram of the human *MNT* locus (not to scale) showing locations of sites (red and blue arrows) for sgRNAs #2 and #4 used in CRISPR/Cas9-mediated gene deletion, and location of the epitope recognised by the MNT antibody. **B,** BL-41 cells have reduced viability after doxycycline induction of *MNT* sgRNA expression. Left panel shows western blot of MNT level at day 4 after treatment with different doses of doxycycline (+Dox), with HSP70 as a loading control. Right panel shows cell viability of control Cas9-infected parental and two sgRNA-infected BL-41 cell lines determined by flow cytometry on day 4 and day 6 after treatment with doxycycline (4 μg/mL). **C**, MNT expression in $Cas9^{+}$ parental and *MNT KO* cell lines, showing that certain *MNT KO* BL-2 clones express smaller MNT polypeptides (see text). **D**, Cytoplasmic localization of truncated MNT proteins in *MNT KO* BL-2 cell lines. Cell lysates were fractionated into nuclear and cytoplasmic fractions (see Methods) and analyzed by western blotting, using HSP70 as a loading control, MAX as a control for a nuclear protein and MCL-1 as a control for a cytoplasmic protein. C=cytosol, N=nucleus. **E**, TaqMan-PCR analysis showing that the increased expression of BIM, PUMA and NOXA proteins in *MNT KO* Rael-BL clones (Fig. 7C) is due, at least in part, to increased transcription of the corresponding genes. Results for independent *MNT KO* lines (colored columns) are expressed as fold-change relative to the corresponding transcripts in the $Cas9^{+}$ parental cell line (black columns) and are plotted as mean of 3 independent experiments +/- SEM;_*P≤ 0.05, **P≤ 0.01, ns= not significant. **F-H** Cell proliferation competition experiments. **F**, Schematic showing parental ($mCherry^{+}GFP^{-}$) Burkitt Lymphoma cells mixed 1:1 with either $mCherry^{+}$ $GFP^{+}$ *MNT KO* clones, or control $MNT^{+/+}$ $mCherry^{+}$ $GFP^{+}$ cells (transfected with sgRNAs but not treated with doxycycline). **G**, Typical flow cytometric analysis of Rael-BL $GFP^{+}$ and $GFP^{-}$ cell populations, showing that GFP-positive *MNT KO* cells are outcompeted by GFP-negative $MNT^{+/+}$ cells (see results in Fig 7F). **H,** Controls showing that GFP-positive $MNT^{+/+}$ cells are not outcompeted by GFP-negative $MNT^{+/+}$ parental cells.
